# Lactate supply overtakes glucose when neural computational and cognitive loads scale up

**DOI:** 10.1101/2022.05.23.493059

**Authors:** Yulia Dembitskaya, Charlotte Piette, Sylvie Perez, Hugues Berry, Pierre J Magistretti, Laurent Venance

## Abstract

The neural computational power is determined by neuroenergetics, but how and which energy substrates are allocated to various forms of memory engram is unclear. To solve this question, we asked whether neuronal fueling by glucose or lactate scales differently upon increasing neural computation and cognitive loads. Here, using electrophysiology, two-photon imaging, cognitive tasks and mathematical modeling, we show that both glucose and lactate are involved in engram formation, with lactate supporting long-term synaptic plasticity evoked by high stimulation load activity patterns and high attentional load in cognitive tasks, and glucose being sufficient for less demanding neural computation and learning tasks. Overall, these results demonstrate that glucose and lactate metabolisms are differentially engaged in neuronal fueling depending on the complexity of the activity-dependent plasticity and behavior.

**One sentence summary:** Neuronal fueling by lactate versus glucose scales differently according to engram level and memory load.

## INTRODUCTION

Brain activity and performance are constrained by the neurovasculature-neuroenergetic coupling (Bullmore and Sporns, 2012; Harris et al., 2012; Li and Sheng, 2022). Neuroenergetics, *i.e*. brain energy metabolism, relies on the blood supply of glucose from the circulation. Blood glucose is taken up during synaptic activity (Attwell et al., 2010; Mann et al., 2021; Tingley et al., 2021), mainly by astrocytes and oligodendrocytes and metabolized into lactate before transport to neurons as energy substrate (Figley and Stroman, 2011; Magistretti and Allaman, 2018; Zimmer et al., 2017; Trevisiol et al., 2017; Pellerin and Magistretti, 1994; Barros and Weber, 2018; Karagiannis et al., 2021; Ivanov et al., 2014; Bonvento and Bolanos, 2021) necessary for optimized coding and memory consolidation (Suzuki et al., 2011; Newman et al., 2011; Alberini et al., 2018; Steadman et al., 2020; Kauffmann et al., 2010; Longden et al., 2014; Murphy-Royal et al., 2020; Zuend et al., 2020; Padamsey et al., 2022). Other fates of glucose include its glial storage in glycogen (Magistretti et al., 1981; Brown and ransom, 2007) to feed the neuronal pentose phosphate pathway (Bolanos et al., 2010; Mergenthaler et al., 2013; Diaz-Garcia et al., 2017; Yellen, 2018) to produce reducing equivalents for long-term memory (de Tredern et al., 2021). Nevertheless, the nature of the energy substrate, glucose *vs* lactate, allocated to various forms of memory engram is not known.

Here, we tested various form of activity-patterns (rate- and time-coding) for Hebbian long-term synaptic plasticity expression in rat CA1 hippocampal pyramidal cells and behavioral tasks with increasing cognitive loads, to determine in which conditions glucose and/or lactate are crucial for engram formation and memory. To this end, using brain slice and *in vivo* electrophysiology, two-photon imaging, mathematical modelling and recognition memory tasks, we show that neuronal lactate is mandatory for demanding neural computation, while glucose is sufficient for lighter forms of activity-dependent long-term potentiation, and that subtle variations of spike amount or frequency are sufficient to shift the energetic dependency from glucose to lactate. Furthermore, we demonstrated that lactate is necessary for a cognitive task requiring high attentional load (object-in-place task) and for the corresponding *in vivo* hippocampal potentiation, but is not needed for a less demanding task (novel object recognition). Our results demonstrate that glucose and lactate metabolisms are differentially engaged in neuronal fueling depending on the complexity of the activity-dependent plasticity and behavior. Beyond reconciling decades-long debate (Barros and Weber, 2018; Diaz-Garcia et al., 2017; Yellen, 2018), our results demonstrate the importance to distinguish specific cellular and molecular mechanisms since the corresponding cognitive perturbations might depend on whether lactate or glucose metabolism is perturbed.

## RESULTS and DISCUSSION

### Rate- and time-coding relied differently on the neuronal lactate

To investigate the relative involvement of glucose and lactate metabolisms in memory formation, we tested two activity-dependent forms of long-term potentiation (LTP) at hippocampal CA1 pyramidal cells (Fig. 1A-C), aiming to reflect two levels of neural computation. We chose two distinct Hebbian activity patterns: (i) a *rate*-coding paradigm involving a high stimulation load (200 stimulations): 5x-theta-burst stimulation (5-TBS), with nested high-frequency (100 Hz) stimulations within slower frequencies (5 Hz) repeated 5 times (at 0.1 Hz), and (ii) a *time*-coding paradigm involving a lower stimulation load (50 stimulations): spike-timing-dependent plasticity (STDP) with 50 pre- and postsynaptic paired stimulations at a low frequency (0.5 Hz). Whole-cell recordings of CA1 pyramidal cells were performed at a physiological glucose concentration (5 mM) (Gruetter et al., 1998), to avoid saturated non-physiological concentrations of glucose (~20-25 mM) classically used in brain slice studies. In the following, drugs were applied intracellularly (i-drug) via the patch-clamp pipette ensuring specific effects in the sole recorded neuron, except in few cases where drugs where bath-applied (e-drug). 5-TBS and STDP paradigms induced LTP (5-TBS-LTP: *p*=0056, n=14; STDP-LTP: *p*=0.0070, n=8), and both were NMDAR-mediated since prevented by the intracellular application of the NMDAR blocker MK801 (i-MK801, 1 mM) (5-TBS: *p*=0.1562, n=7; STDP: *p*=0.6484, n=7) (Fig. 1D and Fig. S1). Since both LTP forms share the same signaling pathway, we could interpret their respective glucose/lactate-dependency based on the activity patterns.

**Figure 1.**
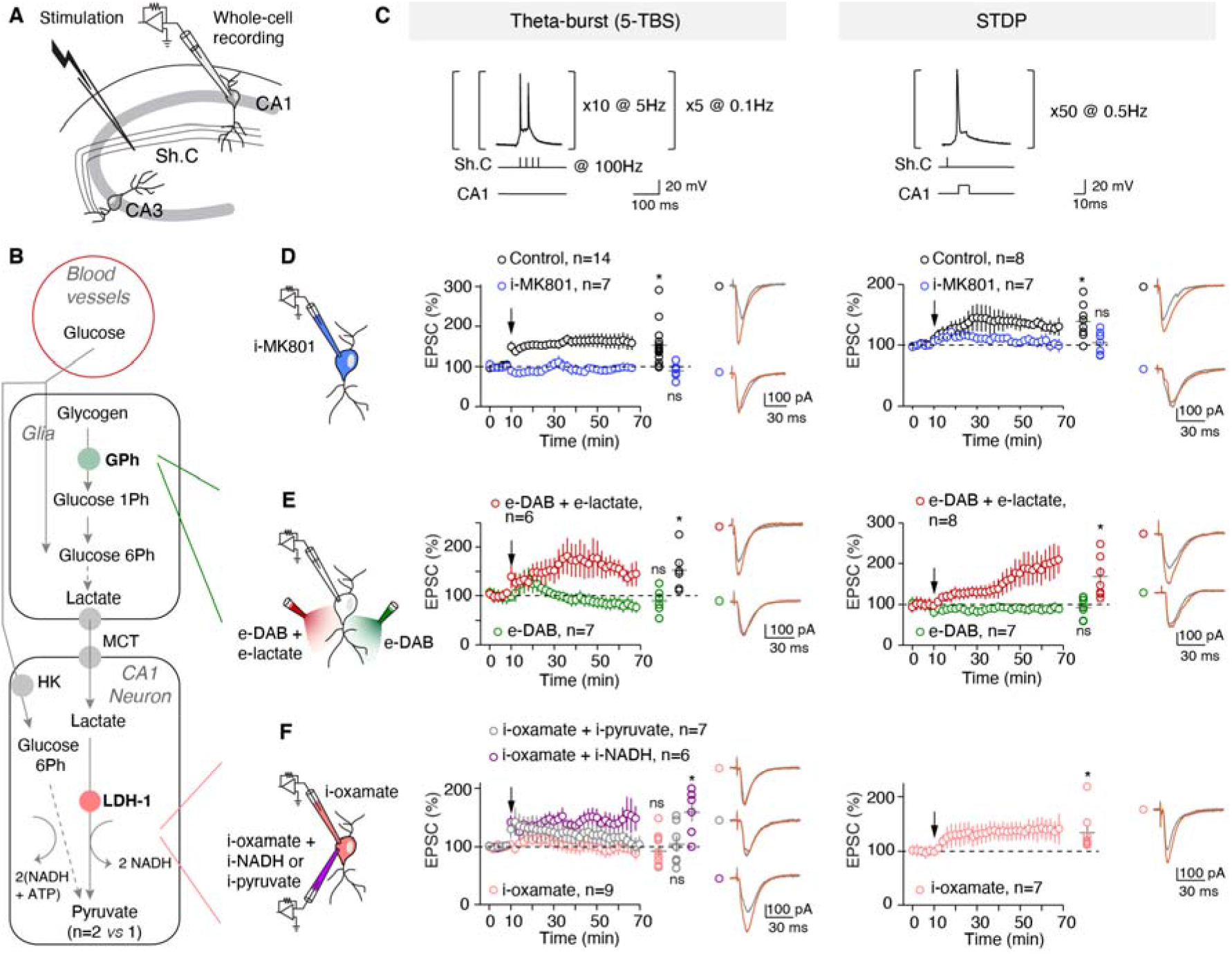
5-TBS-LTP and STDP-LTP relied differently on the neuronal lactate. (**A**) Experimental setup. (**B**) Key steps of the glucose transport and the glia-neuron lactate transport: glycogen catalysis into glucose-1-phosphate via glycogen phosphorylase, lactate entry in neurons via monocarboxylate transporters (MCTs), and lactate conversion to pyruvate by LDH. (**C**) 5-TBS and STDP paradigms. (**D**-**G**) Averaged time-course of the synaptic weight with EPSC amplitude 50-60 min after TBS/STDP. (**D**) 5-TBS-LTP and STDP-LTP were NMDAR-mediated. (**E**) Inhibition of glycogen phosphorylase with e-DAB prevented 5-TBS-LTP and STDP-LTP; Co-application of DAB and lactate allowed 5-TBS-LTP and STDP-LTP. (**F**) Intracellular inhibition of LDH revealed distinct effects on 5-TBS and STDP expression since i-oxamate prevented 5-TBS-LTP but not STDP-LTP. 5-TBS-LTP was rescued with co-application of i-oxamate with i-NADH, but not with i-pyruvate. Representative traces: 15 EPSCs averaged during baseline (grey) and 45 min (red) after protocols (arrows). All data: mean±SEM. *p<0.05; ns: not significant by two tailed *t*-test. See Table S1 for detailed data and statistics.

We first evaluated whether 5-TBS-LTP and STDP-LTP expression equally relies on lactate metabolism (Fig. 1 and Table S1) by sequentially inhibiting two key steps: glycogen mobilization into glucose-1-phosphate, the first step of glycogenolysis leading ultimately to glia-derived lactate, via glycogen phosphorylase, and conversion of lactate into pyruvate via the neuronal lactate dehydrogenase (LDH-1) (Fig. 1B).

We prevented glial glycogenolysis by inhibiting glycogen phosphorylase with 1,4-dideoxy-1,4-imino-D-arabinitol (DAB, 10 μM). With e-DAB, 5-TBS or STDP pairings failed to induce synaptic plasticity (5-TBS: *p*=0.3691, n=6; STDP: *p*=0.4213, n=7) (Fig. 1E). We tested whether lactate overcomes DAB effects. With e-DAB and e-lactate (10 mM), we observed 5-TBS-LTP (*p*=0.0311, n=6) and STDP-LTP (*p*=0.0383, n=7) (Fig. 1E); LTP rescued with e-lactate were not significantly different from control ones (5-TBS-LTP: *p*=0.6735; STDP-LTP: *p*=0.4155). This indicates that lactate formed from glycogenolysis is a key factor for hippocampal LTP induction (Suzuki et al., 2011; Alberini et al., 2018; Murphy-Royal et al., 2020).

We next prevented the conversion of lactate into pyruvate, by applying intracellularly oxamate, an inhibitor of LDH (i-oxamate, 6 mM), only in the recorded CA1 pyramidal cell. Under this condition, 5-TBS did not evoke plasticity (*p*=0.4600, n=9), whereas STDP-LTP could still be observed (*p*=0.0398, n=7) (Fig. 1F). Conversion of lactate into pyruvate was thus required for 5-TBS-LTP but not for STDP-LTP. We then tested which products of lactate conversion by LDH-1, *i.e*. pyruvate or NADH, was needed for 5-TBS-LTP. When i-pyruvate was co-applied intracellularly with i-oxamate, 5-TBS did not induce plasticity (*p*=0.7548, n=7), whereas 5-TBS-LTP was rescued with i-NADH (*p*=0.0145, n=6; vs 5-TBS-LTP control: *p*=0.7995) (Fig. 1F).

As revealed by LDH inhibition, lactate appears as a key element for 5-TBS-LTP (via its metabolism and NADH production), whereas a lighter form of activity-dependent plasticity involving 50 STDP pairings does not depend on the neuronal lactate, and therefore possibly on glucose.

### Confronting mathematical model and experimental data delineates the energetic needs of synaptic plasticity

To provide hypotheses for the differential effects of the neuronal glucose and lactate metabolisms on activity-dependent plasticity, we developed a mathematical model of CA1 synaptic plasticity combined with metabolism. Our model describes the kinetics of a set of signaling and metabolic reactions occurring in a postsynaptic terminal and a nearby interacting glial cell in response to activity patterns (Fig. 2A). The postsynaptic weight is modelled as a bistable system gated by calcium and ATP: postsynaptic calcium triggers LTP when it overcomes a threshold (*LTP*_start_), while the postsynaptic ATP level triggers depotentiation when falling below a second threshold (*ATP*_Thr_). Calcium influx in the postsynaptic neuron changes in response to presynaptic and postsynaptic spikes via the activation of NMDAR and voltage-gated calcium channels. ATP levels in each compartment are computed by a model of metabolic interactions with the astrocyte-neuron lactate shuttle (Jolivet et al., 2015), that includes, among others, glycolysis and LDH activity in glia and the post-synapse, as well as glucose and lactate transfer between them via the extracellular medium. Importantly, the values of the model parameters were estimated using a subset of our experimental data taken from Figures 1D-G and Fig. 2B (*Supplementary Information*, and Tables S1 and S3), while model validation was carried out using model predictions, *i.e*. by checking the accuracy of the model output in experimental conditions that were not used for parameter estimation (the pharmacology perturbation experiments of Fig. 2D-G and Fig. 3). The model captures the amplitude and kinetics of change of the synaptic weight after 5-TBS and STDP pairings (Fig. 2B). In the model, both 5-TBS and STDP paradigms are strong enough to generate large calcium transients in the postsynaptic neuron (Fig. S2A) that overcome the *LTP*_start_ threshold thus triggering LTP. The amplitudes of Na transients in the postsynaptic neuron are much larger with 5-TBS than with STDP, so that ATP consumption by Na,K-ATPases is larger with 5-TBS (Fig. 2C). The availability of lactate as a source of ATP keeps ATP levels well above *ATP*_Thr_ even after 5-TBS. LDH inhibition switches the neuron to a glycolytic regime, an oxidized redox state where ATP level drops to 2.1 mM at rest (Fig. S3). After STDP and with LDH inhibition, the ATP levels keep well above *ATP*_Thr_, while with 5-TBS, ATP quickly fails below *ATP*_Thr_ and the resulting depotentiation forbids LTP expression (Fig. 2C). Experimentally, we found that pyramidal cells recorded with lower i-ATP and i-phosphocreatine (2 and 5 mM, respectively) did not exhibit plasticity following STDP (50 pairings at 0.5 Hz) in control (*p*=0.9439, n=6) or i-oxamate (*p*=0.5992, n=7) (Fig. S4, Table S3A).

**Figure 2:**
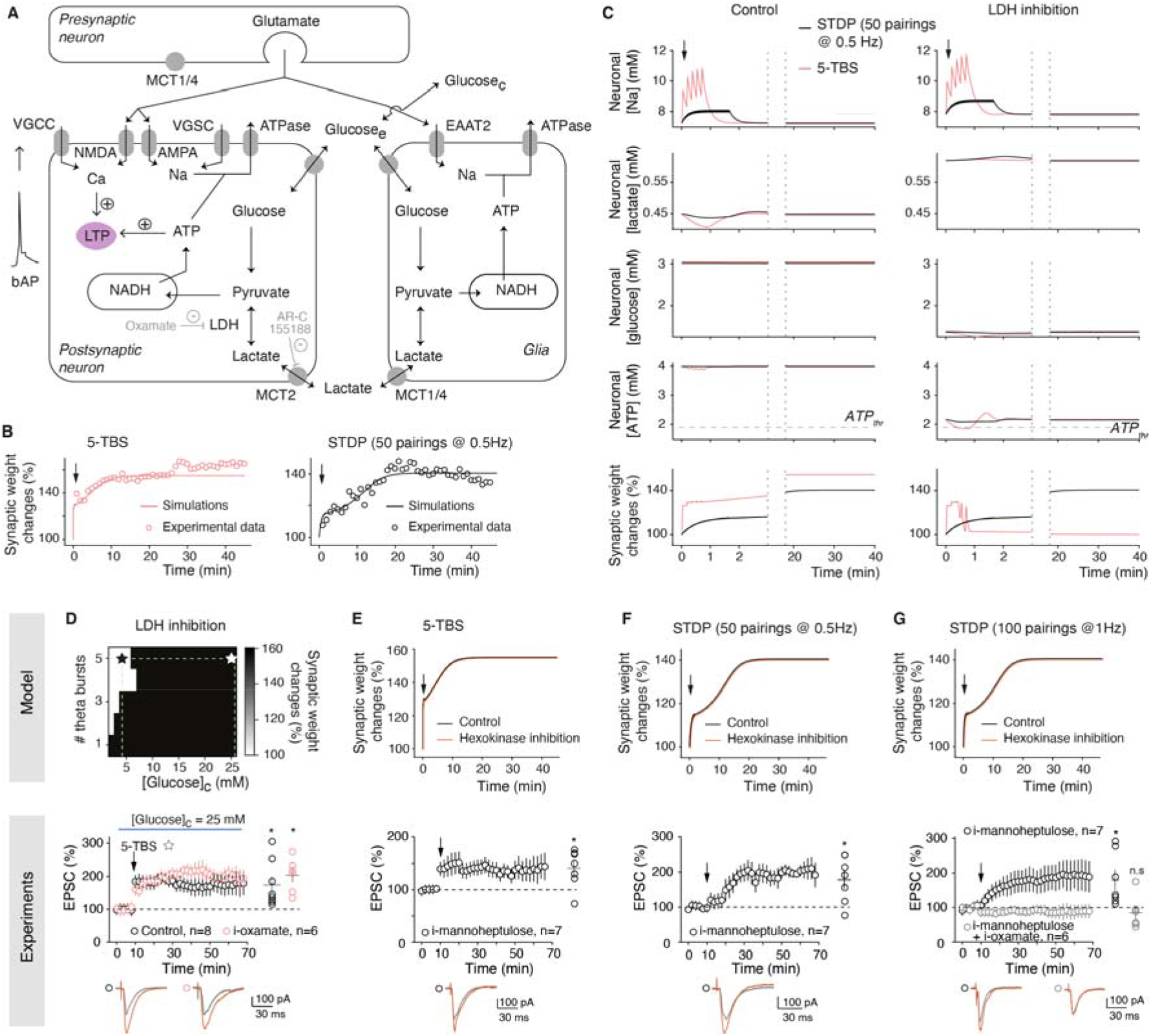
Confronting mathematical model and experimental data delineates the energetic needs of synaptic plasticity. (**A**) In the model (see *Methods, Supplementary Information*, and Table S2), the synaptic weight is gated by both neuronal calcium (potentiation) and ATP (depression); Voltage-gated calcium (VGCC) and sodium (VGSC) channels, EAAT2: excitatory amino acid transporter-2. (**B**) Time course of the synaptic weight in the model (lines), fitted to 5-TBS (n=7) and STDP (n=14; 50 pairings at 0.5Hz, spike-timing=10 ms) experiments (circles). (**C**) Model prediction for the evolution of neuronal concentrations and synaptic weight with 5-TBS (red) or STDP (black). (**D**) TBS-LTP expression depending on glucose concentration ([Glucose]_c_) as predicted by the model. Experimentally 5-TBS induced LTP in high glucose concentration (25 mM), this LTP was not impaired by LDH inhibition (i-oxamate). (**E**) 5-TBS-LTP expression with hexokinase inhibition (i-mannopheptulose). (**F**) 50 pairings at 0.5Hz induced STDP-LTP with i-mannopheptulose. (**G**) 100 pairings at 1Hz induced STDP-LTP with i-mannopheptulose. When neuronal glycolysis and lactate conversion into pyruvate were inhibited with co-applied i-mannopheptulose and i-oxamate, 100 pairings did not induce plasticity. Representative traces: 15 EPSCs averaged during baseline (grey) and 45min (red) after protocols (arrows). All data: mean±SEM (except in 2B where SEM were omitted for clarity). *: p<0.05; ns: not significant by two tailed *t*-test. See Table S3 for detailed data and statistics.

**Figure 3.**
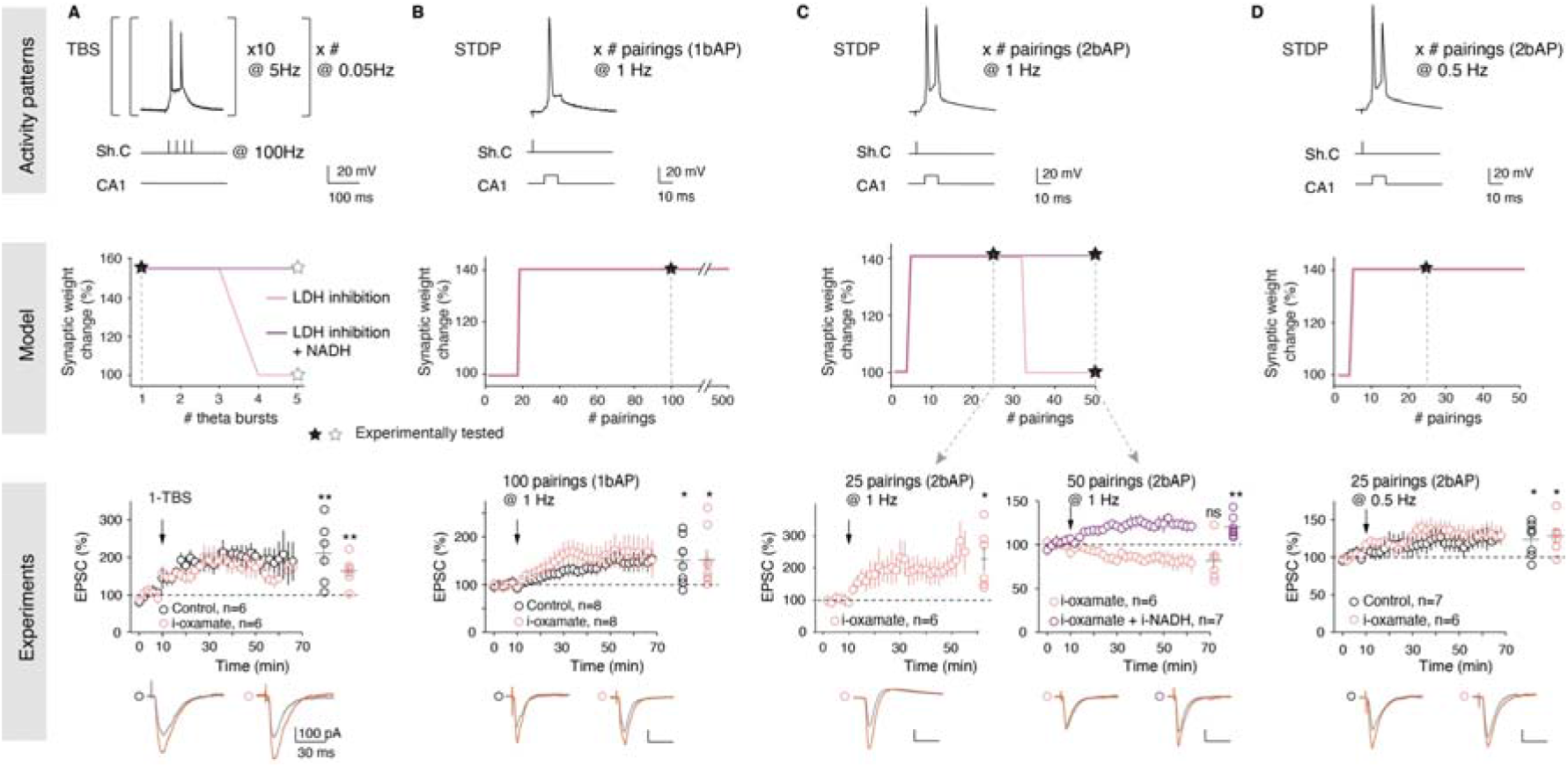
Dependence on lactate for LTP expression is activity-pattern linked. (**A**) Lactate-dependent TBS-LTP depends on the number of TBS (left: protocol). Model predicts that TBS-LTP is dependent on lactate pathway from 2-TBS, as demonstrated experimentally with LTP induced by 1-TBS, which was not impaired by i-oxamate. (**B**) STDP-LTP induced by 20-500 pre-post pairings at 1Hz, with a single bAP/pairing, can be induced under LDH inhibition as predicted by the model and demonstrated experimentally with LTP induced by 100 pairings. (**C**) STDP-LTP induced by pairings at 1Hz with 2 bAPs/pairing, is dependent on LDH activity from 30 pairings as predicted by the model, and demonstrated experimentally with i-oxamate which prevented LTP expression by 50, but not 25, pairings with 2 bAPs; Intracellular co-application of i-oxamate and i-NADH rescued LTP-STDP, as predicted by the model. (**C**) STDP-LTP induced by pairings at 0.5Hz with 2 bAPs/pairings can be induced under LDH inhibition, as predicted by model and demonstrated experimentally with LTP induced by 25 pairings with 2 bAPs/pairings, which is not impaired with i-oxamate. Black stars: experimental conditions tested. Representative traces: 15 EPSCs averaged during baseline (grey) and 45min (red) after protocol (arrows). All data: mean±SEM. *: p<0.05; ns: not significant by two tailed *t*-test. See Table S3 for detailed data and statistics.

Using a model-guided approach (Fig. 2D-G, Table S2), we first investigated the impact of extracellular glucose concentration for 5-TBS-LTP and STDP-LTP expression. Figure 2D shows model output for TBS with LDH inhibited depending on the bath-applied glucose concentration. The main prediction is that LTP should be recovered if bath glucose concentration is large enough. Accordingly, in experiments with high external glucose concentration (25 mM, widely used in brain slices experiments), 5-TBS-LTP (*p*=0.0299, n=8) was no longer sensitive to i-oxamate (*p*=0.0044, n=6) (Fig. 2D), indicating that high glucose concentration bypasses fueling by lactate. Another model prediction is that the inhibition of the hexokinase, the first enzyme of glycolysis catalyzing the phosphorylation of glucose to glucose-6-phosphate (Fig. 1B), should not affect 5-TBS-LTP (Fig. 2E). Experimentally, a specific inhibitor of the hexokinase, mannoheptulose, applied intracellularly (i-mannoheptulose, 10 μM) did not prevent 5-TBS-LTP (*p*=0.0230, n=7) (Fig. 2E), confirming that 5-TBS-LTP relies on lactate and not on glucose metabolism. We next explored the glucose-dependency of STDP, and as predicted by the model, we found that i-mannoheptulose did not prevent STDP-LTP (*p*=0.0210, n=7) (Fig. 2F), indicating that in absence of neuronal glycolysis, the lactate pathway is used for the expression of STDP-LTP (50 pairings at 0.5 Hz). We next doubled the number and frequency of STDP pairings (up to 100 pairings at 1 Hz) to test whether this property could be extended to other STDP forms. In confirmation of the model prediction, we found that LTP induced by 100 pairings at 1 Hz was not impaired i-mannoheptulose (*p*=0.0436, n=6) (Fig. 2G). Interestingly, when both neuronal glucose and lactate sources were impaired by the intracellular co-application of i-mannoheptulose and i-oxamate, (100 pairings at 1 Hz) STDP-LTP was not observed (*p*=0.5297, n=6) (Fig 2G), showing that STDP relies on either glycolysis or lactate pathway.

### Dependence on glucose vs lactate for LTP expression is activity-pattern linked

We varied the TBS/STDP activity patterns to delineate the sensitivity of the plasticity-dependency on glucose and lactate metabolisms. Model calibration relied exclusively on 5-TBS and STDP with 50 pairings at 0.5 Hz (Fig. 2B). We first generated model predictions by varying the number of TBS-bursts and predicted that LTP induction by one single burst (1-TBS) would not be dependent on lactate metabolism, whereas LTP induced with at least 2 bursts (2-TBS) would be, unless performed with high levels of added NADH (Fig. 3A). This was validated experimentally since 1-TBS-LTP (*p*=0.0229, n=6) was not prevented by i-oxamate (*p*=0.0109, n=6) (Fig. 3A). As for predictions related to STDP, we first varied *in silico* the number of pairings and predicted that STDP, even for 500 pairings, would remain non-dependent on lactate metabolism (Fig. 3B). This was demonstrated experimentally since 100 pairings with a single back-propagating action potential (bAP) at 1 Hz induced LTP in control (*p*=0.0191, n=8; Fig. 2G) and with i-oxamate (*p*=0.0391, n=8) with similar magnitude (*p*=0.9836) (Fig. 3B and Table S3). This was also confirmed in mice where i-oxamate prevent 5-TBS-LTP but not LTP induced by STDP with 100 pairings (Fig. S5).

We next varied the number of bAPs per STDP pairings and tested the impact of one additional bAP, *i.e*. going from 1 to 2 bAPs per STDP pairing. Two-photon imaging of dendritic spines and shafts of CA1 pyramidal cells (n=6) showed that the calcium transient triggered by 2 bAPs was roughly twofold compared to 1 bAP (Fig. S6), a feature reproduced by the model (Fig. S2B). For plasticity induction, we kept constant the overall number of postsynaptic stimulations, maintaining a total of 100 pairings (*i.e*. 100 pairings with 1 bAP *vs* 50 pairings with 2 bAPs) and the same frequency (1 Hz). The model predicted that shifting from 1 to 2 bAPs would render STDP lactate-dependent if more than 30 pairings were used (Fig. 3C). Experimentally, we first tested a number of STDP pairings just below the predicted threshold at 30, *i.e*. 25 pairings at 1 Hz with 2 bAPs, and observed that with i-oxamate, LTP was successfully induced (*p*=0.0216, n=6). Furthermore, in full agreement with model prediction, no plasticity was detected for 50 pairings with 2 bAPs (*p*=0.0956, n=6), and adding i-NADH to i-oxamate rescued LTP expression (*p*=0.0061, n=7) (Fig. 3C). These observations illustrate that the on-demand fueling is highly sensitive to the activity patterns on either side of the synapse (Howarth et al., 2012) since a variation from 1 to 2 bAPs was sufficient to render the lactate pathway necessary for LTP expression.

We finally tested whether the number of bAPs *per se* or its combination with the number and frequency of pairings matters. To do so, we kept 2 bAPs per pairings and decreased the number and frequency of pairings two-fold, in order to compare this condition with the STDP paradigm used in Figure 1, *i.e*. 50 postsynaptic stimulations overall (50 pairings with 1 bAP *vs* 25 pairings with 2 bAPs, both at 0.5 Hz) (Fig. 3D and Table S3). The model predicted that in these conditions LTP would be induced and would not depend on lactate. Experimentally, we found that 25 pairings with 2 bAPs (at 0.5 Hz) induced LTP (*p*=0.0369, n=7), and that this LTP was still observed with i-oxamate (*p*=0.0309, n=6) (Fig. 3D).

In conclusion, the dependence on glucose *vs* lactate metabolism precisely scales with the activity patterns used to induce plasticity: neuronal glycolysis is sufficient for light forms of plasticity, and glia-derived lactate is required for more sustained activity-dependent plasticity.

### Inhibition of LDH impaired object-in-place but not novel object recognition learning

Since synaptic plasticity is a major substrate for learning and memory (Nabavi et al., 2014), we next tested whether lactate-dependency scales with learning of recognition memory tasks with increasing cognitive loads. For this purpose, we chose two single-trial tasks involving several structures including hippocampus (Broadbent et al., 2009), which differ by their difficulty level: the object-in-place (OiP) task (with four objects) being more challenging and requiring a higher cognitive load than the novel object recognition (NOR) task (with two objects). These tasks were similarly structured into three phases conducted at one day interval: (1) habituation phase in the empty arena, (2) familiarization phase in presence of two (NOR) or four (OiP) objects, and (3) test in which recognition of new (NOR) or exchanged objects (OiP) was assessed. Rats were injected bilaterally, via cannulas implanted just above CA1 layer, with saline or oxamate (50 mM) solutions 45 min before starting the familiarization phase (Fig. 4).

**Figure 4.**
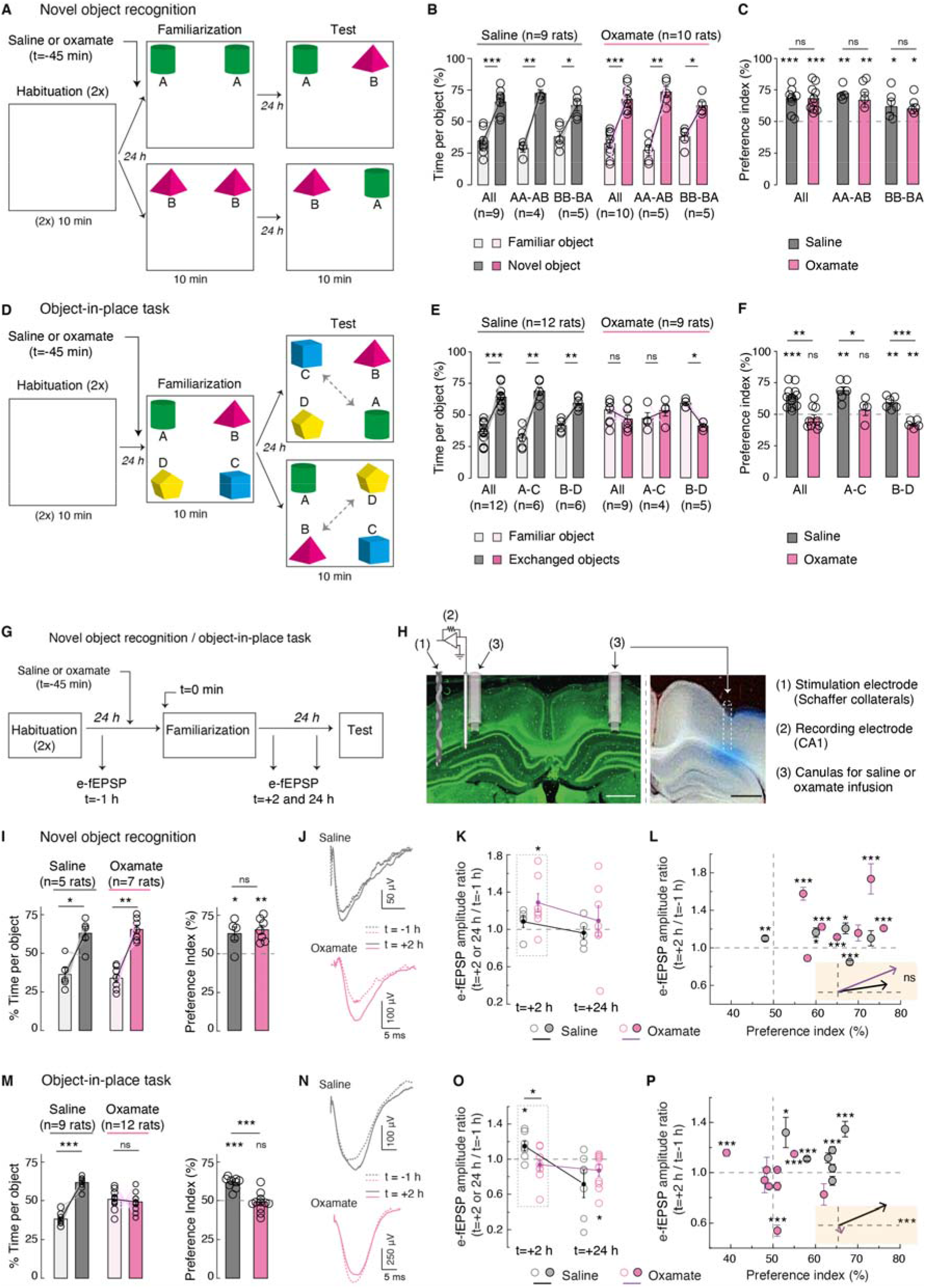
Inhibition of LDH impaired OiP and associated LTP, but not NOR learning. (**A**-**C**) NOR task. (**A**) Experimental set-up. Rats were divided in two subgroups exposed to A-A then A-B, or B-B then B-A, during familiarization and test phases, respectively. (**B** and **C**) Rats injected in CA1 with saline or oxamate (50mM) spent equally more time exploring the novel object. LDH inhibition did not impair novelty detection. (**D**-**F**) OiP task. (**D**) Rats were exposed to A-B-C-D objects during familiarization and were divided in two subgroups experimenting A-C or B-D exchanged-object for test. (**E** and **F**) Saline-injected rats spent more time exploring the exchanged objects, whereas oxamate-injected rats explored equally all objects. LDH inhibition impaired ability to detect place-exchanged objects. (**G**-**P)** e-fEPSP recordings during NOR and OiP tasks. (**G**) Experimental set-up; (**H**) microphotographs showing cannulas and stimulation/recording electrodes locations, and diffusion area (scale bars=1mm). (**I**-**P**) *In vivo* synaptic plasticity during NOR and OiP. e-fEPSPs were recorded before familiarization (baseline) and 2 and 24 hours after familiarization to determine synaptic changes, in relation with behavior. (**I**-**L**) NOR behavioral performance (**I**) with associated LTP of e-fEPSPs after 2h but not after 24h in saline- and oxamate-injected rats (**J**-**L**); averaged vectors show similar trend (**L**). (**M**-**P**) OiP behavioral performance (**M**) with related e-fEPSPs show LTP in saline-but not in oxamate-injected rats 2 hours after familiarization (**N**-**P**); averaged vectors show differences between saline- and oxamate-injected rats (**L**). All data: mean±SEM. *p<0.05; **p<0.01; ***p<0.001; by two-tailed *t*-test. See Tables S4A-D for detailed data and statistics.

For NOR with two objects (A and B), rats injected with saline (n=9) or oxamate (n=10) performed similarly (Fig. 4A-C). Indeed, rats spent more time during the test phase around the new object when considering the time per object and the preference index (saline: *p*=0.0007 and oxamate: *p*=0.0003; saline *vs* oxamate: *p*=0.8126). We ensured that there was no significant preference for object A or B during familiarization rats (Fig. S7A). Furthermore, similar results were found regardless whether the familiarization and test phases were achieved with AA/AB or BB/BA object combination (Fig. 4A-C and Table S4A).

We next subjected rats to the OiP task (Fig. 4D-F): rats were left for familiarization with four different objects (A, B, C and D) for which they did not show preference (Fig. S7B)), and then for the test phase two of the four objects were place-exchanged (from ABCD to CBAD or ADCB).We found that rats injected with saline (n=12) spent more time around the exchanged objects, whereas rats injected with oxamate (n=9) did not notice the exchange since they continued exploring equally the four objects as shown by the time per object (Fig. 4E) and the preference index (Fig. 4F) (saline: *p*<0.0001 and oxamate: *p*=0.2500; saline *vs* oxamate: *p*=0.0002). (Fig. 4D-F and Table S4B). Similar results were found regardless whether the familiarization and test phases were performed with ABCD/CBAD or ABCD/ADCB object exchange combination.

In conclusion, when conversion of lactate into pyruvate is impaired before familiarization, rats still succeeded in NOR, but not in a more challenging task, OiP.

### Inhibition of LDH impaired OiP-induced LTP, but not NOR-induced LTP

We further examined in NOR and OiP tasks whether synaptic plasticity occurred with CA1 hippocampal infusion of saline or oxamate in relation with behavioral performance (Fig. 4G-P, Fig. S7C-G and Tables S4A-D). Synaptic weights were evaluated *in vivo* by monitoring evoked-field-EPSPs (e-fEPSPs) at synapses between Schaffer collaterals and CA1 pyramidal cells in behaving rats. To do so, we placed chronic stimulating and recording electrodes in Schaffer collaterals and CA1, respectively, in rats equipped bilaterally with cannulas for saline or oxamate infusion (Fig. 5B). We first ensured that this set of rats performed in NOR and OiP similarly as aforementioned (Fig. 4I and 4M, Table S4A-B). To determine plasticity expression, e-fEPSPs were monitored before (t=-1 hour, before saline or oxamate injection) and after (t=+2 and +24 hours) the familiarization phase (Fig. 4J-L and 4N-P, Fig. S7C-G, Table S4C-D).

In the NOR task, LTP of e-fEPSPs dominates at t=+2 hours in both saline- and oxamate-injected rats (n=5 and 7, respectively), and is followed by a scaled reduction of synaptic weights at t=+24 hours (as shown by the positive correlation between plasticities at t=+2 and +24 hours), leading on average to no plasticity at t=+24 hours (Fig. 4J and 4L, Fig. S7D and Table S4C). The behavioral and plasticity profiles were similar between saline- and oxamate-injected rats, as illustrated by the average vectors (*p*=0.412; insert in Fig. 4L) considering the preference index and plasticity at t=+2 hours after familiarization (Fig. 4L, Fig. S7F and Table S4D).

A different picture was obtained for OiP: e-fEPSPs exhibited LTP at t=+2 hours in saline-injected rats, but not in oxamate-injected rats (n=9 and 12, respectively) (Fig. 4O). More precisely, when considering e-fEPSPs plasticity at +2 hours in relation with behavioral performance, all saline-injected rats detected exchanged objects and 4/7 displayed LTP, whereas among the 78% (7/9) of the oxamate-injected rats that did not detect the exchange, only one showed LTP while the others exhibited an absence of plasticity or long-term depression (LTD). This is also illustrated by the difference between averaged vectors (*p*<0.001; insert in Fig. 4P, and Table S4D). Monitoring of the synaptic weights 24 hours after familiarization, showed similar plasticity pictures for saline- and oxamate-injected rats, *i.e*. LTD or absence of plasticity, despite distinct preference indexes (Fig. S7E and S7G and Table S4D).

In conclusion, rats detecting novelty in the OiP task displayed LTP after the familiarization phase, whereas oxamate-injected rats which were not able to detect novelty, did not show LTP. Therefore, learning novelty in a challenging memory task (OiP) requires lactate-dependent LTP while glucose-dependent LTP can be sufficient to learn a less demanding cognitive task (NOR).

Here, we show that scaling of the computational and cognitive loads requires the metabolism of glycogen-derived lactate to match the energetic requirements of sustained neural activity patterns and high cognitive load. For less demanding plasticity and learning paradigms, glucose suffices as an energy substrate. We thus reconcile conflicting views concerning the involvement of lactate *vs* glucose in synaptic plasticity (Barros and Weber, 2018; Diaz-Garcia et al., 2017; Yellen, 2018; Bak and Walls, 2018). The two pools of energy substrates (glucose and lactate) can be distinctly allocated on-demand (Kasischke et al., 2004; Chuquet et al., 2010; Mächler et al., 2016; Ruminot et al., 2017; Hall et al., 2012) in qualitative (activity hotspots) and quantitative (engram levels) manners, within the hard limit of the global energy availability of cellular metabolism (Clarke and Sokoloff, 1999; Bruckmaier et al., 2020). We delineated the domains of activity pattern for which LTP expression requires glucose and/or lactate metabolism, and their borders defined by the elementary elements of neural computation, *i.e*. the rate and timing codes. This is particularly illustrated by the fact that variation of a single bAP was sufficient to shift the LTP-dependency from glucose to lactate. The glia-derived lactate, as well as neuronal glycolysis, could thus be triggered after extracellular potassium changes as low as ~200 μM, according to theoretical estimations of the potassium efflux upon single AP (Saetra et al., 2021), consistent with the demonstrated glycogenolytic action of K^+^ (Hof et al., 1988; Bittner et al., 2011; Choi et al., 2012). The demonstration that under conditions of glycogenolysis inhibition, lactate, but not glucose, allows sustained electrical activity (Trevisiol et al., 2017), fear or spatial learning (Suzuki et al., 2011; Newman et al., 2011; Alberini et al., 2018) (that involves high perceptual load) is in line with our results. Also, a decrease in lactate production mediated by mitochondrial cannabinoid type-1 receptor activation in astrocytes, alters social behavior in a lactate reversible manner (Jimenez-Blasco et al., 2020). In contrast fast learning engrams originating from light activity patterns (Piette et al., 2020), could emerge even in the absence of lactate metabolism, with glucose as the main energy substrate.

The tight dependence and sensitivity of neuronal signaling on energy availability renders the brain vulnerable to conditions in which energy delivery or utilization are compromised. This is the case for neurodegenerative diseases such as Alzheimer’s and Parkinson’s diseases, amyotrophic lateral sclerosis and frontotemporal dementia (Morrison et al., 2012; Camandola and Mattson, 2017; Cunnane et al., 2020; Pathak et al., 2013; Bonvento and Bolanos, 2021) as well as for neurodevelopmental disorders such as glucose transporter-1 deficiency syndrome (Tang et al., 2019). Pharmacological strategies aimed at boosting brain energy metabolism by acting at specific cellular and molecular targets (*e.g*. neurons vs glial cells, glycolysis *vs* glycogenolysis, molecular steps of the ANLS) deserve close attention as they may provide an original and unifying interventional approach for diseases characterized by cognitive impairment and neurodegeneration.

## Acknowledgments

This work was supported by fundings from Collège de France, INSERM, CNRS and Fondation Bettencourt Schueller. Y.D. was supported by ANR Dopaciumcity, C.P. by Ecole Normale Supérieure.

We thank the Venance lab members for helpful suggestions and critical comments. We thank Giuseppe Gangarossa and Marika Nosten-Bertrand for their helpful suggestions for behavioral tasks; Marie Vandecasteele for technical and analysis assistance for preliminary *in vivo* electrophysiological recordings; Ilya Prokin for the custom-made software for calcium transient analysis.

## Authors contributions

Conceptualization: LV, PJM and HB

Experimental investigation: YD, CP and SP

Mathematical modeling: HB.

Supervision: LV, PJM

Writing: LV, PJM and HB

Funding acquisition: LV

## Methods

### Animals

Experiments were conducted in male Sprague Dawley rats (Charles River, L’Arbresle, France) P_30-35 days_ for brain slice patch-clamp and two-photon imaging, and P_7-9 weeks_ (weight: 250-300 g) for behavioral tasks and *in vivo* electrophysiology. In a subset of experiments, C57BL/6 mice P_28-35_ days were used for brain slice electrophysiology (Fig. S5). All experimental protocols and methods were approved by the local animal welfare committee (Center for Interdisciplinary Research in Biology Ethics Committee) and EU guidelines (directive 2010/63/EU). Every effort was made to minimize animal suffering and to use the minimum number of animals per group and experiment. Animals were housed in standard 12-hour light/dark cycles and food and water were available *ad libitum*.

### Patch-clamp whole-cell recordings in brain slices

Transverse hippocampal slices (350μm-thick) were prepared using a vibrating blade microtome (7000smz-2, Campden Instruments Ltd., UK) in ice-cold cutting solution containing (in mM): 93 N-Methyl-D-glucamine-Cl, 2.5 KCl, 30 NaHCO_3_, 1.2 NaH_2_PO4, 20 HEPES, 5 Na-ascorbate, 0.5 CaCl_2_, 1 MgSO_4_*7H_2_O, 25 glucose, 3 Na pyruvate. The slices were transferred to the storage solution containing (in mM): 125 NaCl, 2.5 KCl, 5 glucose, 25 NaHCO_3_, 1.25 NaH_2_PO_4_, 2 CaCl_2_, 1 MgCl_2_ with 10 μM pyruvic acid, for one hour at 34°C and then to room temperature. In a subset of experiments (Fig. 2D) storage solution containing 25 mM of glucose was used, as specified. All solutions were saturated with 95% O_2_ and 5% CO_2_.

For whole-cell recordings from CA1 pyramidal neurons, borosilicate glass pipettes of 3-5 MΩ resistance were filled with (in mM): 127 K-gluconate, 30 KCl, 10 HEPES, 10 phosphocreatine (or 5 as specified in Fig. S4), 4 Mg-ATP (or 2 as specified in Fig. S4), 0.3 Na-GTP, 0.1 EGTA (adjusted to pH 7.35 with KOH). The composition of the extracellular solution was (mM): 125 NaCl, 2.5 KCl, 5 (or 25 as specified in Fig. 2D) glucose, 25 NaHCO_3_, 1.25 NaH_2_PO_4_, 2 CaCl_2_, 1 MgCl_2_ and 10 μM pyruvic acid, through which 95% O_2_ and 5% CO_2_ was bubbled. Signals were amplified with EPC10-2 amplifiers (HEKA Elektronik, Lambrecht, Germany). Current- and voltage-clamp recordings were sampled at 20 kHz, with the Patchmaster v2×32 program (HEKA Elektronik). All recordings were performed at 35°C, using a temperature control system (Bath-controller V, Luigs & Neumann, Ratingen, Germany) and slices were continuously perfused with extracellular solution, at a rate of 2 ml/min.

### Synaptic plasticity induction protocols

Synaptic responses in CA1 pyramidal cells were evoked by electrical stimulations of Schaeffer’s collaterals with concentric bipolar electrodes (Phymep, Paris, France) placed in *stratum radiatum* area of hippocampus, at a distance exceeding 200 μm from the recording site. Electrical stimulations were monophasic, at constant current (ISO-Flex stimulator, AMPI, Jerusalem, Israel). Currents were adjusted to evoke 100-300 pA EPSCs. Repetitive control stimuli were applied at 0.1 Hz. Recordings were made over a period of 10 minutes at baseline, and for at least 60 minutes after the synaptic plasticity induction protocols; long-term changes of synaptic weight were measured in the last 10 minutes. Experiments were excluded if the mean input and series resistance (Ri and Rs, respectively) varied by more than 20% through the experiment.

#### Theta-burst stimulation (TBS)

TBS of the Schaeffer’s collaterals consisted of 10 bursts (single burst: 4 stimuli of 0.2 ms duration at 100 Hz) repeated at 5 Hz. TBS were applied either as a single TBS (1-TBS; Fig. 3A) or repeated 5 times at 0.1 Hz (5-TBS).

#### Spike-timing-dependent plasticity (STDP) protocols

STDP protocols consisted of pairings of pre- and postsynaptic stimulations separated by a specific and fixed time interval (ΔtSTDP=+10-15 ms, *i.e*. presynaptic stimulation preceded postsynaptic activation) (Feldman, 2012). Presynaptic stimulations corresponded to Schaeffer’s collaterals stimulations and the postsynaptic stimulation of an action potential (AP) evoked by a depolarizing current step (10 ms duration) in the recorded CA1 pyramidal cell. In a subset of experiments (Fig. 3C and 3D), 2 postsynaptic APs were elicited, as specified. Paired stimulations were repeated *n* times at a *f* frequency. In Figures 1 and S1, we used STDP with 50 pairings at 0.5 Hz. In Figure 2 and 3, the number of postsynaptic APs (1 or 2), number of pairings (25, 50 or 100) and frequency (0.5 or 1 Hz) were varied, as specified.

### Patch-clamp data analysis

Off-line analysis was performed with Fitmaster (Heka Elektronik), Igor-Pro 6.0.3 (Wavemetrics, Lake Oswego, OR, USA) and custom-made software in Python 3.0. Statistical analysis was performed with Prism 5.02 software (San Diego, CA, USA). We individually measured and averaged 60 successive EPSCs, comparing the last 10 minutes of the recording with the 10-minute baseline recording in each condition using t-test. In all cases “n” refers to a single cell experiment from a single brain slice. All results are expressed as mean±SEM. Statistical significance was assessed by two-tailed student t-tests (unpaired or paired t-tests) or one-way ANOVA (with Newman-Keuls post hoc test) when appropriate, using the indicated significance threshold (*p*).

### Two-photon imaging combined with whole-cell patch-clamp recordings

Morphological tracer Alexa Fluor 594 (50 μM) (Invitrogen, Waltham, MA, USA) and calcium-sensitive dye Fluo-4F (250 μM) (Invitrogen) were added to the intracellular solution to monitor calcium transients. Cells were visually identified under Scientifica TriM-Scope II system (LaVision, Germany), with a 60x/1.00 water-immersion objective. Alexa Fluor 594 and Fluo-4F were excited at 830 nm wavelength (femtoseconds IR laser Chameleon MRU-X1; Coherent, UK), and their fluorescence were detected with photomultipliers within 525/50 and 620/60 nm ranges, respectively. Line-scan imaging at 200 Hz was performed to obtain calcium signals in the dendritic shaft and spines and was synchronized with patch-clamp recordings. In each recording, we produced somatic depolarization and monitored maximal Ca2+ elevations to verify linear dependence of Fluo-4F Ca2+ signals (nonlinearity was below 20%) and that Ca2+ transients were below saturation level. The changes in baseline Ca2+ level were monitored as the ratio between the baseline Fluo-4F and Alexa Fluor 594 fluorescence. The cell was discarded for ratio>20%. The dark noise of the photomultipliers was collected when the laser shutter was closed in every recording. One or two back-propagating action potentials (bAPs) evoked by a depolarizing current step (10 ms duration) were applied to monitor dendritic calcium transients. Electrophysiological data and calcium transients were analyzed with Fitmaster (Heka Elektronik) and/or custom-made software in Python 3.0 (https://github.com/calciumfilow/calcium-trace-extractor). The measurements of calcium transient were represented as ΔG/R: (G_peak_-G_baseline_)/(R_baseline_-R_dark noise_). Baseline Ca2+ signals were represented by baseline G/R, (G_baseline_-G_dark noise_)/(R_baseline_-R_dark noise_), where G is the Fluo-4F fluorescence, and R is Alexa Fluor 594 fluorescence. G_baseline_, R_baseline_ and G_peak_ were obtained from the parameters of the bi-exponential fitting model in each trial and then averaged between 5-6 repetitions for each condition; the bi-exponential fitting was chosen to maximize the fit of the calcium-evoked events and gave better fit quality than single- and triple-exponential fitting (https://github.com/calciumfilow/calcium-trace-extractor). G_dark noise_ and R_dark noise_ are the dark currents of the corresponding photomultipliers. We ensured that R_baseline_ and G_baseline_/R_baseline_ ratio did not exceed 20% over recording. The statistical significance was tested using a paired or unpaired Student’s t-test in Prism 5.02 software (San Diego, CA, USA).

### Chemicals

1,4-Dideoxy-1,4-imino-D-arabinitol hydrochloride (DAB) (10μM) and DAB+L-lactate (10mM) were dissolved directly in the extracellular solution. (+)-5-methyl-10,11-dihydroxy-5H-dibenzo(a,d)cyclohepten-5,10-imine (MK801) (1mM), Na-oxamate (6mM), Na-oxamate (6mM) + L-pyruvate (10mM), Na-oxamate (6mM) + K_2_NADH (4mM), D-Mannoheptulose (10μM), D-Mannoheptulose (10μM) + Na-oxamate (6mM), were added to intracellular recording solution. Na-oxamate (50mM) was dissolved directly in the saline solution for *in vivo* experiments. All chemicals were purchased from Sigma (Saint-Quentin Fallavier, France) except for MK801 and AR-C155858 (Tocris, Ellisville, MO, USA).

For patch-clamp experiments, drugs were applied intracellularly (noted: i-drugs), ensuring specific intracellular effect in the sole recorded neuron without affecting the neighboring neurons or astrocytes; in few cases (Fig. 1E), drugs were applied extracellularly (noted: edrugs).

### Cannulas implants and *in vivo* microinjections

Under anesthesia with pentobarbital sodium (50 mg/kg i.p.), rats (250–300 g) were placed in a stereotactic apparatus, the cranium was exposed and two holes were drilled to aseptically implant double guide cannulas (26GA, Bilaney, Germany) into dorsal hippocampi bilaterally (stereotaxic coordinates: anteroposterior =-4.0mm from bregma; mediolateral =−2.5mm from midline). The guide cannulas were secured with dental cement. The two internal cannulas (33GA, Bilaney, Germany) were then inserted within the guide cannulas (dorsoventral =-2.2mm from bone surface). After surgery, 1mg/g i.p. of analgesic agent (Metacam, 1.5mg/ml solution) was administered for 3 days. Rats were allowed to recover from surgery for 7 days before behavioral test beginning. Location of the cannulas are illustrated in Figure 4H.

Rats were injected bilaterally through the stainless-steel cannulas 45 min before the familiarization phase of behavioral testing (novelty-object recognition or object-in-place). Cannulas were briefly connected to 10 μL Hamilton syringes by means of polyethylene tubes. Rats were injected with either saline (0.8 mM sterile NaCl) or with Na-oxamate (50 mM) diluted in saline; 1-2 μl per hippocampus using automatic infusion pump at a speed of 0.2 μl/min.

### Procedures for novel object recognition (NOR) and object-in-place (OiP) tasks

NOR and OiP tasks were run in a black Plexiglas training arena of squared shape (100×100cm with 50cm walls). The arena was placed in a sound-attenuated room with a controlled light intensity of 50 lx. The objects used were made out of plastic (laboratory cylinders and Lego blocks) and were selected to induce comparable attraction (Fig. S7A-B). The objects were fixed on the arena floor with a 10 cm distance from the walls. The NOR (2 objects: A and B) and OiP (4 objects: A, B, C and D) tasks involved three sessions upon three consecutive days: habituation (on day 1), familiarization (on day 2) and test (on day 3).

#### Habituation (NOR and OiP)

Habituation phase was similarly conducted for the NOR and OiP tasks. The first day, rats were introduced into the empty arena (in its center) for 10 min to habituate to the arena; this procedure was repeated twice the first day with 4 h intersession interval.

#### NOR task

For the familiarization session on day 2, the rat was placed in the center of the arena and exposed to two identical objects for 10 min: two objects A for half of the animals and two objects B for another half of the rats. Then, rats were returned to their home cage. For the test session (or retrieval) on day 3, the rat was placed back into the center of the arena for object discrimination and were exposed to one familiar object and a novel test object, B for group precedingly exposed to A-A and A for group exposed to B-B, for 10 min.

#### OiP task

For the familiarization session on day 2, the rat was placed in the center of the arena and exposed to four different objects: A, B, C, D, located clockwise as A-B-C-D, for 10 min. For the test session on day 3, the rat was placed back into the center of the arena for object discrimination and were exposed to the exchanged position of two of the presented objects, C-B-A-D for half of the animals and A-D-C-B for another half, for 10 min.

#### NOR and OiP performance analysis

Animals were videotaped during familiarization and test sessions. Videos were analyzed and time spent on exploration each object (sniffing or licking) was measured during familiarization and test sessions. For assessing NOR and OiP memory performance, the calculation of object discrimination, the exploration time of the novel object was expressed as percentage of the total exploration time of familiar and novel objects during familiarization and test sessions. Then, we calculated the relative time of exploration per object and the preference index (%) in control (saline-injected rats) and in oxamate (Na-oxamate-injected rats).

### *In vivo* electrophysiology in behaving rats during object-in-place task

Field excitatory postsynaptic potentials (fEPSP) evoked from Schaffer collaterals stimulation (e-fEPSP) were measured in the left CA1 over the 3-day behavioral assessment in rats subjected to NOR (familiarization: A-A and test: A-B) or OiP (familiarization: A-B-C-D and test: C-B-A-D) tasks. A recording wire (stainless steel, 0.005” diameter) was stereotaxically implanted under pentobarbital-ketamine anesthesia (pentobarbital: 30 mg/kg i.p., Ceva Santé Animale, Libourne, France; ketamine: 27.5 mg/kg, i.m., Imalgène, Mérial, Lyon, France) in CA1 stratum radiatum along the guide cannula (anteroposterior: −4.1mm, mediolateral: 2.5, dorsoventral : 3.3), and a bipolar stimulating electrode (2 twisted wires, same as the recording electrode) in the ipsilateral Schaffer collaterals (anteroposterior: −4.4mm, mediolateral: 4.4, dorsoventral : 3.4). The rat body temperature was maintained using at 37°C during surgery. Rats were allowed at least a week of recovery before electrophysiological recordings and behavioral testing. Recordings were performed during 10-15 min in a 24×44cm Plexiglas box, 60 min before familiarization (before the oxamate or saline injection) and, 2 and 24 hours after familiarization. Schaffer e-fEPSPs were amplified, and acquired at 20 kHz using a KJE-1001 system (Amplipex, Szeged, Hungary). Test pulses (150-200 μs duration, 20-450 μA) were evoked every 30 seconds using a square pulse stimulator and stimulus isolator (Model 2100, AM-Systems, Sequim, WA, USA). Responses were analyzed offline using custom Matlab codes (2019b, The Mathworks, Natick, MA, USA) scripts. Traces was smoothed (linear averaging) over 0.15, 0.25 or 0.5 ms depending on the noise level. Detection of local extrema was performed to define the start and peak of the response, and to measure the e-fEPSP amplitude. Statistical analyses (two-tailed Student t-test) and calculation of plasticity ratio were made on average on 25 trials. The average baseline of e-fEPSP amplitude obtained at t=-60 min was used to normalize every individual e-fEPSP amplitude at +2 and +24 hours, which were then averaged to obtain the plasticity ratio. The F-statistic and *p*-value associated with the average vectors were obtained from MANCOVA test (two dependent variables: preference index and plasticity ratio; one factor: belonging to the saline- or oxamate-injected rat group). Pearson’s correlation coefficients were computed for paired plasticity ratios measured at 2 or 24 hours after familiarization phase.

### Histology

After the completion of NOR and OiP experiments, cannulas and recording and stimulation electrodes positions were examined. Rats were anaesthetized (sodium pentobarbital, 150mg/kg i.p.) and transcardially perfused with saline followed by 4% ice-cold paraformaldehyde in 0.1 M phosphate buffer. Brains were extracted, post-fixed in PFA (4% in PBS) for 2 days at 4°C and then cryoprotected in 30% sucrose solution for one week. Brains were cut in coronal 50 μm sections using a cryotome (HM400, Microm Microtech, Francheville, France) and slices were maintained in 0.1 M potassium-PBS (pH=7.4). The sections were Nissl-stained with thionin (Neurotrace 500/525, Thermofischer, Waltham, MA, USA), and images were acquired using a stereozoom fluorescence microscope (Axiozoom, Zeiss, Oberkochen, Germany) and processed in ImageJ. Only the rats with cannula tips located bilaterally within the dorsal hippocampi were included in the final data analysis.

### Mathematical model

In our experiments, we did not differentiate between lactate coming from astrocytes or oligodendrocytes. In the mathematical model, though, parameter calibration imposes to specify which cell type one considers. The vast majority of published models and data on the subject is specific of astrocytes; we opted for astrocytes as glial cells in the model. Our model therefore simulates the network of signaling and metabolic reactions occurring in a postsynaptic neuronal terminal and an interacting astrocyte shown in Figure 2A.

#### Stimulations

The membrane potential of the postsynaptic compartment *V_n_* is given by:

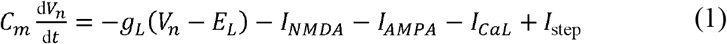

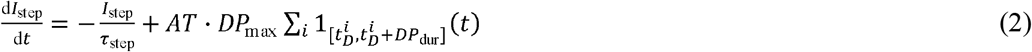

with the indicator function 1_[*a,b*]_(*x*) = 1 if *x* ∈ [*a, b*], 0 otherwise, 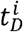 is the time of the beginning of *i*^th^ depolarization of the postsynaptic neuron, *DP*_max_ is the amplitude of the depolarization current in the postsynaptic neuron and *DP*_dur_ its duration. *τ*_step_ is the time step of the step current, *AT* is the attenuation of the bAP at the postsynaptic compartment and the models for the ionic currents *I_NMDA_,I_AMPA_,I_CaL_* (Fig. 2A) are given in the Supplementary Information.

##### STDP 1bAP

For STDP protocols with a single bAP per postsynaptic stimulation, we emulated the bAP by incrementing the postsynaptic potential by a constant value, *i.e*.:

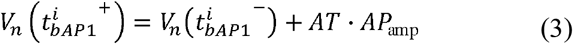

where 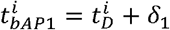 is the time of the (first) bAP, peaking with a delay *δ*_1_ after the beginning of the depolarization and *AP*_amp_ is the amplitude of the bAP measured in the soma.

##### STDP 2bAP

For STDP protocols with two bAPs per postsynaptic stimulation, we added an additional bAP at time 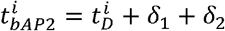 where *δ*_2_ is the delay between the two bAPs:

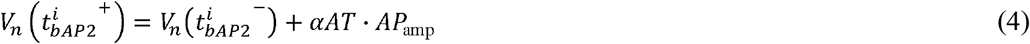

where *α* is the attenuation of the second bAP compared to the first bAP. During STDP protocol, we first set the times of each presynaptic stimulation 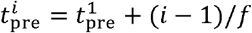 where 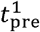 is the time of the first presynaptic stimulation (arbitrary) and *f* is the stimulation frequency in Hz. The postsynaptic times are then set according to the spike timing 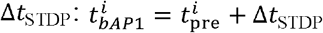 and 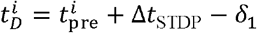

##### TBS

one presynaptic theta burst is composed of 4 presynaptic stimulations at 100 Hz repeated 10 times at 5 Hz and the frequency of the theta-burst themselves is 0.1 Hz, *i.e*.:
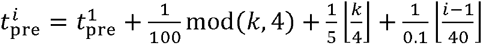

where ⌊*x*⌋ denotes the integer part of *x*, mod(*x, j*) = *x* − *j*⌊*x/j*⌋ is its modulo *j* and *k* = mod(*i* − 1,40). On the postsynaptic side, experimental observations show that during a presynaptic TBS, the probability for a presynaptic spike to trigger a postsynaptic bAP is circa 0.5, therefore we set *DP*_max_ = 0 (no postsynaptic current injected in TBS) and set *TP*_max_ = 0 for odd *i*s.

#### STDP

In the postsynaptic compartment, the variation of cytosolic calcium concentration is computed as:

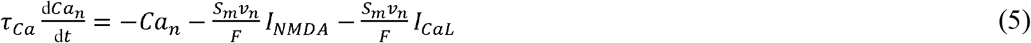

where *S_m_v_n_* is the surface-to-volume ratio of the postsynaptic compartment and *F* the Faraday constant. In the model, both the cytosolic calcium concentration and the cytosolic ATP concentration drive a bistable internal signaling state summarized by the state variable *ρ* ∈ [0,1]^53^:

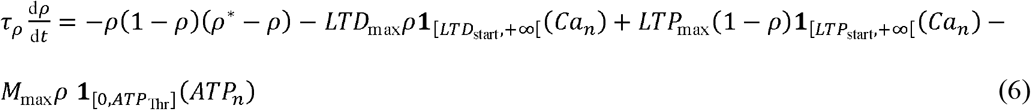

where *ATP_n_* is the concentration of ATP in the postsynaptic compartment, *LTD*_max_, *LTP*_max_ and *M*_max_ are, respectively, the amplitudes of the calcium-gated LTD and LTP and of the ATP-gated depotentiation and *LTD*_start_, *LTP*_start_ and *ATP*_Thr_. their respective thresholds. *ρ\** sets the value of the unstable (intermediate) steady-state of the bistable and *τ_ρ_* the time scale to reach the two stables steady-states *ρ* = 0 and *ρ* = 1. Finally, the synaptic weight is taken an affine function of the signaling state variable:

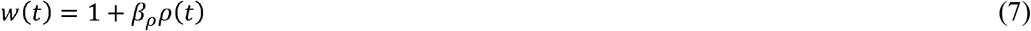

#### Astrocyte-neuron lactate shuttle

To derive the time course of *ATP_n_* in eq. (6) as a function of the pre- and post-synaptic stimulations, we used the model developed by Jolivet *et al* (2015). In both the postsynaptic terminal and a nearby interacting astrocyte (Fig. 2A), the model accounts for glycolysis, *i.e*. the production of pyruvate from glucose with glyceraldehyde-3 phosphate (GAP) and phosphoenolpyruvate (PEP) as intermediates as well as pyruvate formation from lactate by lactate dehydrogenase (LDH). The model also incorporates the production of NADH from pyruvate by the TCA cycle in the mitochondria, resulting ATP production by the electron transport chain as well as NADH shuttling from the cytosol to the mitochondria. Glucose and lactate are exchanged between the cytosol of the two compartments and the periplasmic extracellular medium via transporters, including MCT2 for lactate transport from the periplasmic volume to the postsynaptic compartment. Diffusive transport also occurs between the periplasmic volume and reservoir solutions with fixed concentrations (bath solution or blood capillaries of the slices). In addition, the model accounts for the activity-dependent dynamics of sodium ions in both compartments. Presynaptic stimulations trigger Na influx via EEAT2 channels in the astrocyte and voltagegated sodium channels (VGSC) and (partly) AMPA receptors in the postsynaptic compartment. Sodium is then pumped back by Na,K-ATPases (ATPase), consuming ATP in the process.

We have implemented the model of Jolivet *et al* (2015) *in extenso*. Our only change has been to substitute the Hodgkin-Huxley equation used for the postsynaptic membrane voltage in Jolivet *et al* (2015) by our eq. (1). This made us slightly adapt the maximum conductance for the postsynaptic VGSC, the strength of the effect of the presynaptic stimulation on the astrocyte and the leak conductance. All the remaining of the model, *i.e*. the 28 ODEs and the 80+ remaining parameters were taken unchanged from Jolivet *et al* (2015).

#### Parameter estimation

In total, coupling the two models (STDP+astrocyte-neuron lactate shuttle) left us with 27 parameters to estimate (Table S2). To this end, we used a subset of our available experimental data (training set), whereas validation was carried out by checking the accuracy of the model output for experimental conditions that were not used in parameter estimation (pharmacology perturbation experiments, changes of extracellular glucose concentrations, different stimulation protocols). See Supplementary Information for a complete description of the model as well as parameter estimation strategy and values.

## Data availability

the data that support the findings of this study are available from the corresponding author upon reasonable request.

## Code availability

the computer code for the model is available online open-source at https://gitlab.inria.fr/hberry/anls_stdp

