## Supplemental Information for "Lactate supply overtakes glucose when neural computational and cognitive loads scale up"

**This PDF file includes:**

Supplementary Methods

References for the Supplementary Methods

Supplementary Figures 1 to 7 and corresponding Legends

Supplementary Tables 1 to 4 and corresponding Legends

**Supplementary Methods**

**Mathematical model**

**Ionic currents of the postsynaptic compartment membrane.** The AMPA current was modeled as a double exponential conductance (Roberts, 2014):

$I_{AMPA}=\left[ g_{AMPA\text{max}} \sum_{i} \left( \text{exp}\left( -\frac{t-t_{\mathrm{pre}}^{i}}{\tau_{AMPA1}} \right)-\text{exp}\left( -\frac{t-t_{\mathrm{pre}}^{i}}{\tau_{AMPA2}} \right) \right)\mathbf{1}_{\left[ t_{\mathrm{pre}}^{i},+\infty\right[}\left( t \right) \right]\left( V_{n}-E_{AMPA} \right)$	(S1)

where $t_{\mathrm{pre}}^{i}$ is the time of the *i*^th^ presynaptic stimulation, $g_{AMPA\text{max}}$ the maximal AMPA conductance and $\tau_{AMPA1}$ and $\tau_{AMPA2}$ the two-time scales. The NMDAR was also modelled with a double exponential^55^:

$I_{NMDA}=\left[ g_{NMDA\text{max}} B\sum_{i} \left( \text{exp}\left( -\frac{t-t_{\mathrm{pre}}^{i}}{\tau_{NMDA1}} \right)-\text{exp}\left( -\frac{t-t_{\mathrm{pre}}^{i}}{\tau_{NMDA2}} \right) \right)\mathbf{1}_{\left[ t_{\mathrm{pre}}^{i},+\infty\right[}\left( t \right) \right]\left( V_{n}-E_{NMDA} \right)$	(S2)

where the magnesium block $B=\left( 1+\left[ \text{Mg}^{2+} \right]/3.57\text{exp}\left( -0.062V_{n} \right) \right)^{-1}$. The VGCC current was modelled after (Solinas et al., 2014) as a high-voltage L-type current:

$I_{CaL}=g_{CaL\text{max}} s^{2}u\left( V_{n}-E_{Ca} \right)$	(S3)

with activation and inactivation functions given by:

$\frac{\text{d}\text{s}}{\text{d}t}=50\left( 1-s \right)\text{exp}\left( \frac{V_{n}+29.06}{15.9} \right)-80s \text{exp}\left( \frac{{-V}_{n}-18.66}{25.6} \right)$	(S4)

and

$\frac{\text{d}\text{u}}{\text{d}t}=\left( 1-u \right)\text{exp}\left( \frac{{-V}_{n}-48}{18.2} \right)-u\text{exp}\left( \frac{V_{n}+48}{83} \right)$	(S5)

The voltage-gated sodium current was taken from Jolivet *et al* (2015):

$I_{Na}=g_{Na\text{max}} {m_{\infty}}^{3}h\left( V_{n}-E_{Nan} \right)$	(S6)

where the steady-state activation was $m_{\infty}={\alpha_{m}}/\left( \alpha_{m}+\beta_{m} \right)$ with $\alpha_{m}=-0.1\left( V_{n}+33 \right)/\left[ \text{exp}\left( -0.1\left( V_{n}+33 \right) \right)-1 \right]$ and $\beta_{m}=4\text{exp}\left( -\left( V_{n}+58 \right)/{12} \right)$ and the inactivation was given by:

$\frac{\text{d}\text{h}}{\text{d}t}=0.07\left( 1-h \right)\text{exp}\left( \frac{-V_{n}-50}{10} \right)-\frac{h}{\text{exp}\left( \frac{{-V}_{n}-20}{10} \right)+1}$	(S7)

Note that $I_{Na}$ does not contribute to the evolution of the membrane potential in eq.(1) since eq.(1) does not include the spike currents but emulates the bAP using a simple increment of the potential at the spiking time followed by an exponential decay.

**Astrocyte-neuron lactate shuttle.** We implemented the model of Jolivet *et al* (2015) where $S_{x}$ denotes the concentration of chemical species S in compartment $x=\left\{ a,n \right\}$ for astrocyte or postsynaptic neuronal, respectively. We give below a complete depiction of the equations and parameters of the model, but interested readers should refer to the original paper for further specifics.

The cytosolic sodium concentration is given by

$\frac{\text{d}{Na}_{x}}{\text{d}t}=\frac{S_{m}v_{x}}{F}g_{Nax}\left( E_{Nax}-V_{x} \right)-3S_{m}v_{x}k_{\text{pump}x}{Na}_{x}\frac{{ATP}_{x}}{1+{{ATP}_{x}}/{K_{\text{Mpump}}}}+J_{\text{stim}x}$	(S8)

with the Nernst potential $E_{Nax}={RT}/F\text{log}\left( {Na}_{e}/{Na}_{x} \right)$, $g_{Nax}$ the leak conductance and $k_{\text{pump}x}$ the maximal rate of the Na-K-ATPase pump in compartment *x*. $J_{\text{stim}x}$ is the sum of sodium influxes triggered by the stimulations. In the postsynaptic compartment:

$J_{\text{stim}n}=\frac{S_{m}v_{x}}{F}\left( -\frac{2}{3}I_{AMPA}-I_{Na} \right)$	(S9)

while in the astrocyte:

$J_{\text{stim}a}=2.25\times{10}^{-5}\times1500f\mathbf{1}_{\left[ t_{\mathrm{pre}}^{1},t_{\mathrm{pre}}^{N} \right]}\left( t \right)$	(S10)

where *N* is the total number of presynaptic stimulations and *f* their frequency (Jolivet *et al*., 2015).

Cytosolic glucose concentrations are given by

$\frac{\text{d}{GLC}_{n}}{\text{d}t}={Tg}_{en}\left( \frac{{GLC}_{e}}{{GLC}_{e}+K_{tg}}-\frac{{GLC}_{n}}{{GLC}_{n}+K_{tg}} \right)-k_{HKPFKn}\frac{{ATP}_{n}}{1+\left( {{ATP}_{n}}/{K_{I,ATP}} \right)^{4}}\frac{{GLC}_{n}}{{GLC}_{n}+K_{g}}$	(S11)

and

$\frac{\text{d}{GLC}_{a}}{\text{d}t}={{Tg}_{ca}\left( \frac{{GLC}_{c}}{{GLC}_{c}+K_{tg}}-\frac{{GLC}_{a}}{{GLC}_{a}+K_{tg}} \right)+Tg}_{ea}\left( \frac{{GLC}_{e}}{{GLC}_{e}+K_{tg}}-\frac{{GLC}_{a}}{{GLC}_{a}+K_{tg}} \right)-k_{HKPFKa}\frac{{ATP}_{a}}{{1+\left( {{ATP}_{a}}/{K_{I,ATP}} \right)}^{4}}\frac{{GLC}_{a}}{{GLC}_{a}+K_{g}}$	(S12)

where ${GLC}_{e}$ is the extracellular concentration of glucose, i.e. in the pericellular volume, while ${GLC}_{c}$ is its – constant - concentration in the large (reservoir) volume of the bath solution and/or the blood vessels. ${Tg}_{xy}$ is the constant of glucose transport between compartments *x* and *y,* $k_{HKPFKx}$ is the maximal rate for the part of glycolysis from hexokinase to phosphofructokinase (lumped as a single equivalent reaction).

Glyceraldehyde-3 phosphate (GAP) concentrations are obtained with:

$\frac{\text{d}{GAP}_{x}}{\text{d}t}=2k_{HKPFKx}\frac{{ATP}_{x}}{{1+\left( {{ATP}_{x}}/{K_{I,ATP}} \right)}^{4}}\frac{{GLC}_{x}}{{GLC}_{x}+K_{g}}-k_{PGKx}{GAP}_{x}{ADP}_{x}\frac{N-{NADH}_{x}^{c}}{{NADH}_{x}^{c}}$	(S13)

where $k_{PGKx}$ is the maximal rate for the part of glycolysis from GAP dehydrogenase to enolase, ${NADH}_{x}^{c}$ is the concentration of NADH in the cytosol of compartment *x* and *N* is the total concentration of NADH, i.e. the sum of NADH and NAD+ concentration in the cytosol. Phosphoenolpyruvate (PEP) concentration obeys:

$\frac{\text{d}{PEP}_{x}}{\text{d}t}=k_{PGKx}{GAP}_{x}{ADP}_{x}\frac{N-{NADH}_{x}^{c}}{{NADH}_{x}^{c}}-k_{PKx}{PEP}_{x}{ADP}_{x}$	(S14)

with $k_{PKx}$ the pyruvate kinase rate in *x*. Next, the dynamics of pyruvate concentration (PYR) is given by:

$\frac{\text{d}{PYR}_{x}}{\text{d}t}=k_{PKx}{PEP}_{x}{ADP}_{x}-k_{LDH\text{on}x}{PYR}_{x}{NADH}_{x}^{c}+k_{LDH\text{off}x}{LAC}_{x}\left( N-{NADH}_{x}^{c} \right)-v_{\text{mitoin}x}\frac{{PYR}_{x}}{K_{Mmito}+{PYR}_{x}}\frac{{N-NADH}_{x}^{m}}{N-{NADH}_{x}^{m}+K_{MNADx}}$	(S15)

Here $k_{LDH\text{on}x}$ and $k_{LDH\text{off}x}$ are the forward and reverse rates, respectively, of lactate dehydrogenase (LDH) in compartment *x*, ${NADH}_{x}^{m}$ is the concentration of NADH in the mitochondrion of *x* and $v_{\text{mitoin}x}$ is the maximal rate of the TCA cycle. Lactate (LAC) dynamics is obtained via:

$\frac{\text{d}{LAC}_{n}}{\text{d}t}=k_{LDH\text{on}n}{PYR}_{n}{NADH}_{n}^{c}-k_{LDH\text{off}n}{LAC}_{n}\left( N-{NADH}_{n}^{c} \right)-{Tl}_{ne}\left( \frac{{LAC}_{n}}{{LAC}_{n}+K_{tlne}}-\frac{{LAC}_{e}}{{LAC}_{e}+K_{tlne}} \right)$	(S16)

where ${Tl}_{xy}$ is the constant of lactate transport between compartments *x* and *y* and ${LAC}_{e}$ is the extracellular concentration of lactate, *i.e.* in the pericellular volume. Likewise, in the astrocyte:

$\frac{\text{d}{LAC}_{a}}{\text{d}t}=k_{LDH\text{on}a}{PYR}_{a}{NADH}_{a}^{c}-k_{LDH\text{off}a}{LAC}_{a}\left( N-{NADH}_{a}^{c} \right)-{Tl}_{ae}\left( \frac{{LAC}_{a}}{{LAC}_{a}+K_{tlae}}-\frac{{LAC}_{e}}{{LAC}_{e}+K_{tlae}} \right)-{Tl}_{ac}\left( \frac{{LAC}_{a}}{{LAC}_{a}+K_{tlac}}-\frac{{LAC}_{c}}{{LAC}_{c}+K_{tlac}} \right)$	(S17)

where ${Tl}_{xy}$ is the constant of lactate transport between compartments *x* and *y* and ${LAC}_{c}$ is the extracellular concentration of lactate in the reservoir volume. Likewise, the concentration of lactose in the pericellular extracellular medium is given by:

$\frac{\text{d}{LAC}_{e}}{\text{d}t}=\frac{{Tl}_{ae}}{r_{ea}}\left( \frac{{LAC}_{a}}{{LAC}_{a}+K_{tlae}}-\frac{{LAC}_{e}}{{LAC}_{e}+K_{tlae}} \right)+\frac{{Tl}_{ne}}{r_{en}}\left( \frac{{LAC}_{n}}{{LAC}_{n}+K_{tlne}}-\frac{{LAC}_{e}}{{LAC}_{e}+K_{tlne}} \right)-{Tl}_{ec}\left( \frac{{LAC}_{e}}{{LAC}_{e}+K_{tlec}}-\frac{{LAC}_{c}}{{LAC}_{c}+K_{tlec}} \right)$	(S18)

and that of glucose:

$\frac{\text{d}{GLC}_{e}}{\text{d}t}=-\frac{{Tg}_{ea}}{r_{ea}}\left( \frac{{GLC}_{e}}{{GLC}_{e}+K_{tg}}-\frac{{GLC}_{a}}{{GLC}_{a}+K_{tg}} \right)-\frac{{Tg}_{en}}{r_{en}}\left( \frac{{GLC}_{e}}{{GLC}_{e}+K_{tg}}-\frac{{GLC}_{n}}{{GLC}_{n}+K_{tg}} \right)+{Tg}_{ce}\left( \frac{{GLC}_{c}}{{GLC}_{e}+K_{tg}}-\frac{{GLC}_{e}}{{GLC}_{e}+K_{tg}} \right)$	(S19)

with $r_{xy}$ the ratio between the volume of compartment *x* and that of compartment *y*. The concentration of cytosolic NADH (${NADH}_{x}^{c}$) is obtained by integration of:

$\left( 1-\xi\right)\frac{\text{d}{NADH}_{x}^{c}}{\text{d}t}={k_{PGKx}{GAP}_{x}{ADP}_{x}\frac{N-{NADH}_{x}^{c}}{{NADH}_{x}^{c}}-k}_{LDH\text{on}x}{PYR}_{x}{NADH}_{x}^{c}+k_{LDH\text{off}x}{LAC}_{x}\left( N-{NADH}_{x}^{c} \right)-T_{NADHx}\frac{{R^{-}}_{x}}{{R^{-}}_{x}+M_{cytox}}\frac{{R^{+}}_{x}}{{R^{+}}_{x}+M_{mitox}}$	(S20)

where $T_{NADHx}$ is the maximal rate for NADH shuttling from cytosol to the mitochondria sub-compartment, $\xi$ is the relative mitochondria volume, ${R^{-}}_{x}=\frac{{NADH}_{x}^{c}}{N-{NADH}_{x}^{c}}$ and ${R^{+}}_{x}=\frac{N-{NADH}_{x}^{m}}{{NADH}_{x}^{m}}$. For the mitochondrial concentration of NADH, one gets:

$\xi\frac{\text{d}{NADH}_{x}^{m}}{\text{d}t}=T_{NADHx}\frac{{R^{-}}_{x}}{{R^{-}}_{x}+M_{cytox}}\frac{{R^{+}}_{x}}{{R^{+}}_{x}+M_{mitox}}+4v_{\text{mitoin}x}\frac{{PYR}_{x}}{K_{Mmito}+{PYR}_{x}}\frac{{N-NADH}_{x}^{m}}{N-{NADH}_{x}^{m}+K_{MNADx}}-v_{\text{mitoout}x}\frac{{O_{2}}_{x}}{K_{O2mito}+{O_{2}}_{x}}\frac{{ADP}_{x}}{{ADP}_{x}+K_{MADPx}}\frac{{NADH}_{x}^{m}}{{NADH}_{x}^{m}+K_{MNADHx}}$	(S21)

where $v_{\text{mitoout}x}$ is the maximal rate of the electron transport chain in *x* and ${O_{2}}_{x}$the oxygen concentration in this compartment. ATP concentration in the postsynaptic compartment is given by:

$\left( 1-\frac{{dAMP}_{n}}{{dATP}_{n}} \right)\frac{\text{d}{ATP}_{n}}{\text{d}t}=-2k_{HKPFKn}\frac{{ATP}_{n}}{{1+\left( {{ATP}_{n}}/{K_{I,ATP}} \right)}^{4}}\frac{{GLC}_{n}}{{GLC}_{n}+K_{g}}+k_{PGKn}{GAP}_{n}{ADP}_{n}\frac{N-{NADH}_{n}^{c}}{{NADH}_{n}^{c}}+k_{PKn}{PEP}_{n}{ADP}_{n}-J_{ATPasesn}-S_{m}V_{n}k_{\text{pump}n}{Na}_{n}\frac{{ATP}_{n}}{1+{{ATP}_{n}}/{K_{\text{Mpump}}}}+3.6v_{\text{mitoout}n}\frac{{O_{2}}_{n}}{K_{O2mito}+{O_{2}}_{n}}\frac{{ADP}_{n}}{{ADP}_{n}+K_{MADPn}}\frac{{NADH}_{n}^{m}}{{NADH}_{n}^{m}+K_{MNADHn}}+k_{\text{CKon}n}{PCr}_{n}{ADP}_{n}-k_{\text{CKoff}n}\left( C-{PCr}_{n} \right){ATP}_{n}$

(S22)

with $J_{ATPasesn}$ a parameter accounting for ATPase activities outside Na-K-ATPases, $k_{\text{CKon}n}$ and $k_{\text{CKoff}n}$ the forward and backward rates of creatine kinase, respectively, and *C* the total concentration of creatine plus phosphocreatine.

${{dAMP}_{n}}/{{dATP}_{n}}$, the ratio between deoxyAMP and deoxyATP is computed with ${{dAMP}_{x}}/{{dATP}_{x}}=-1+0.5q_{AK}-0.5\sqrt{u_{x}}+{Aq_{AK}}/\left( {ATP}_{x}\sqrt{u_{x}} \right)$ where $q_{AK}$ is the adenylate kinase equilibrium constant, $A={AMP}_{x}+{ADP}_{x}+{ATP}_{x}$ is the total adenine nucleotide concentration and $u_{x}={q_{AK}}^{2}+4q_{AK}\left( A/{{ATP}_{x}}-1 \right)$. Similarly, the concentration of ADP is computed from that of ATP using ${ADP}_{x}={0.5ATP}_{x}\left( -q_{AK}+\sqrt{u_{x}} \right)$.

Now, in the astrocyte, the concentration of ATP is given by:

$\left( 1-\frac{{dAMP}_{a}}{{dATP}_{a}} \right)\frac{\text{d}{ATP}_{a}}{\text{d}t}=-2k_{HKPFKa}\frac{{ATP}_{a}}{{1+\left( {{ATP}_{a}}/{K_{I,ATP}} \right)}^{4}}\frac{{GLC}_{a}}{{GLC}_{a}+K_{g}}+k_{PGKa}{GAP}_{a}{ADP}_{a}\frac{N-{NADH}_{a}^{c}}{{NADH}_{a}^{c}}+k_{PKa}{PEP}_{a}{ADP}_{a}-J_{ATPasesa}-{\frac{3}{4}J}_{pumpa0}-{\frac{7}{4}S}_{m}V_{a}k_{\text{pump}a}{Na}_{a}\frac{{ATP}_{a}}{1+{{ATP}_{a}}/{K_{\text{Mpump}}}}+3.6v_{\text{mitoout}a}\frac{{O_{2}}_{a}}{K_{O2mito}+{O_{2}}_{a}}\frac{{ADP}_{a}}{{ADP}_{a}+K_{MADPa}}\frac{{NADH}_{a}^{m}}{{NADH}_{a}^{m}+K_{MNADHa}}+k_{\text{CKon}a}{PCr}_{a}{ADP}_{a}-k_{\text{CKoff}a}\left( C-{PCr}_{a} \right){ATP}_{a}$

(S23)

where phosphocreatine concentrations are given by

$\frac{\text{d}{PCr}_{x}}{\text{d}t}=- k_{\text{CKon}x}{PCr}_{x}{ADP}_{x}+k_{\text{CKoff}x}\left( C-{PCr}_{x} \right){ATP}_{x}$	(S24)

Finally, oxygen concentrations are obtained through:

$\frac{\text{d}{0_{2}}_{x}}{\text{d}t}=\frac{{PS}_{cap}}{v_{x}}\left( K_{O2}\left( \frac{Hb.OP}{{0_{2}}_{c}}-1 \right)^{-1/n_{h}}-{0_{2}}_{x} \right)-0.6v_{\text{mitoout}x}\frac{{O_{2}}_{x}}{K_{O2mito}+{O_{2}}_{x}}\frac{{ADP}_{x}}{{ADP}_{x}+K_{MADPx}}\frac{{NADH}_{x}^{m}}{{NADH}_{x}^{m}+K_{MNADHx}}$	(S25)

where ${{PS}_{cap}}/{v_{x}}$ is the oxygen transport constant from the reservoir to the cells and ${0_{2}}_{c}$ oxygen concentration in the reservoir.

**Numerical integration.** The system of ODEs consisting of eq.(1)-(7) and eq.(S1)-(S25) was integrated using a variable-order adaptive-stepsize stiff solver (ode15s in Matlab®), interrupting integration at every stimulation events (times$t_{D}^{i},t_{D}^{i}+{DP}_{\text{dur}}$, $t_{bAP1}^{i}$and $t_{bAP2}^{i}$ if relevant) to account for the discontinuities of eq.(2-4). The system was first integrated for 2x10^5^ seconds in the absence of any electrical stimulation to guaranty that the initial state (before electrical stimulation) is the stable steady-state. After electrical simulation (TBS or STDP), the system was integrated for 45 minutes. The state of the system 45 min after the stimulation sets the value of the synaptic weight (eq.7) after the stimulation, thus the potential expression of LTP.

**Parameter estimation.** To calibrate the model, we used the following strategy. All the parameters of equations (S3) to (S25) kept the values set by Jolivet *et al* (2015) (as indicated in Table S2) except for $g_{Na\text{max}}$ (eq. S6), $g_{\text{L}}$(eq.1) and the prefactor of eq.(S10), that were modified as explained below. Therefore, parameter estimation was almost completely restricted to the parameters of eq.(1-7). Together, this represented 27 parameters to estimate (Table S2). Parameter values were estimated on a subset of our available experimental data (see below) whereas validation was carried out by checking the accuracy of the model output for experimental conditions that were not used in parameter estimations.

1. The parameters of the postsynaptic stimulations ${DP}_{\text{max}}$, $\tau_{\text{step}}$, ${AP}_{\text{amp}}$ and $\alpha$as well as the delays $\delta_{1}$and $\delta_{2}$ and the leak conductance $g_{\text{L}}$were fitted to the experimental traces of the postsynaptic membrane potential as measured in the soma by the patch electrode (Fig. S2B). The attenuation factor of the bAP between the soma and the postsynaptic compartment, *AT*, was set to a value that roughly corresponds to a synapse located at mid-distance between the soma and the dendrite end (Migliore et al., 2005) (see their Fig 4D).
2. $g_{AMPA\text{max}}$, $\tau_{AMPA1}$ and $\tau_{AMPA2}$ were set to yield EPSPs of 2 mV amplitude, with short onset and a decay time scale around 10 ms, as measured in dendrites (Watanabe et al., 2002). $\tau_{NMDA1}$ and $\tau_{NMDA2}$ were set to yield a NMDA-component for the calcium influx that rises fast and takes roughly 200 ms to get back to zero and $g_{NMDA\text{max}}$ was fixed so that each presynaptic spike increases the cytosolic calcium level by 0.17 mM at -70 mV (Sabatini et al., 2002). $g_{CaL\text{max}}$was set so that one bAP (on top of the depolarizing current) triggers a calcium influx of 400 nM amplitude at the synapse in the absence of presynaptic stimulation (Carter and Sabatini, 2004). We also checked that with these parameter values, the amplitude of the calcium peak triggered by a postsynaptic stimulation comprising two bAPs is indeed roughly twice the amplitude obtained with a single bAP (Fig. S2B) as measured in our experimental setup both in spines and shafts (Fig. S6).
3. In the original model ^(^Jolivet *et al*., 2015), the membrane voltage of the presynaptic compartment is modelled as a Hodgkin-Huxley equation with variable Nernst potentials for Na. Because of the variable Nernst potential, this model exhibits strong spike-frequency adaptation so the postsynaptic neuron quickly ceases to emit spikes after the onset of the stimulation. Since our membrane voltage for the postsynaptic compartment does not exhibit spike-frequency adaptation, we had to adapt a pair of parameters to guaranty that the electrical stimulations employed in the experiments used to calibrate the model by Jolivet *et al* (2015 (their Figure 4A, with experimental data taken from^33^ will still yield the correct time course in our model. We therefore adapted the values of $g_{Na\text{max}}$ and the prefactor of eq.(S10) so that the stimulation used in Kasischke et al. (2004) (presynaptic stimulations for 20 seconds) yielded in our model the same time-courses for${NADH}_{a}^{c}$ and ${NADH}_{n}^{m}$as their experimental measurements (their Figure 4D). In particular (Fig. S2C) this stimulation leads in our model to 1) a dip of ${NADH}_{n}^{m}$ of around -10%, peaking around the end of the effective stimulation, and converging back to baseline after 15-20 min, and 2) a delayed overshoot of ${NADH}_{a}^{c}$ that peaks at approx. +8% roughly 20 min after the end of the stimulation. Those dynamics reproduce previous experimental measurements (Kasischke et al., 2004), in both quantitative and qualitative terms.
4. The parameters related to the dynamics of the synaptic weight were set as follows. The calcium thresholds ${LTD}_{\text{start}}$ and ${LTP}_{\text{start}}$ were set so that a “standard” STDP protocol (1 bAP at 1 Hz) triggers LTP for positive spike timings that are not larger than 30 ms and for more than approx. 20 pairings (Fig. S2D). The values of ${LTD}_{\text{max}}$, ${LTP}_{\text{max}}$, $\rho^{*}$and $\beta_{\rho}$ were fitted on the experimental measurements of the time course of the synaptic weight change for a STDP protocol with 1 bAP, 50 x at 0.5 Hz (Fig. 2B). Note that with 5-TBS, the experimental measurement of the final amplitude of the synaptic weight change was larger than with STDP (Fig. 2B), we therefore adapted the value of $\beta_{\rho}$ for 5-TBS (but kept the values of ${LTD}_{\text{max}}$, ${LTP}_{\text{max}}$ and $\rho^{*}$ to those obtained by fitting on STDP 0.5 Hz 50x).
5. Finally, we fixed the ATP threshold ${ATP}_{\text{Thr}}$to a value that allows discriminating between STDP 0.5 Hz 50 pairings (LTP) stimulations and 5-TBS with oxamate (no LTP). In particular, the astrocyte-neuron lactate shuttle model predicts that neuronal ATP, ${ATP}_{n}$, falls below 2 mM for 5-TBS in the presence of oxamate (Fig. 2C), but remains well above 2 mM for 5-TBS in control conditions and STDP 0.5 Hz 50 pairings (in control and with oxamate). In the absence of experimental data to set the value of $M_{\text{max}}$, we used a value large enough to cancel LTP for 5-TBS in the presence of oxamate.

**Simulation of pharmacological experiments.** The action of pharmacological agents was emulated by changing the corresponding parameters to the following values:

1. *Oxamate*: the forward and backward rate constants of LDH in the postsynaptic neuronal compartment were divided tenfold, i.e. we set$k_{LDH\text{on}n}=0.723$ mM^-1^s^-1^ and $k_{LDH\text{off}n}=0.0720$mM^-1^s^-1^.
2. *Mannoheptulose:* the forward constant of the HKPFK reaction in the postsynaptic neuronal compartment was divided by 1000, *i.e.* we set $k_{HKPFKn}=0.0504\times{10}^{-3}$ mM^-1^s^-1^
3. *Changes of glucose concentration in the bath:* was emulated by a corresponding change of the glucose concentration in the (constant) reservoir, ${GLC}_{c}$.
4. *Changes of NADH in the patch pipette:* were emulated by corresponding change of the total NADH+NAD^+^ concentration *N* in the postsynaptic compartment (while *N* kept its control value of 0.212 mM in the astrocyte).

**Supplementary Figures and Legends**

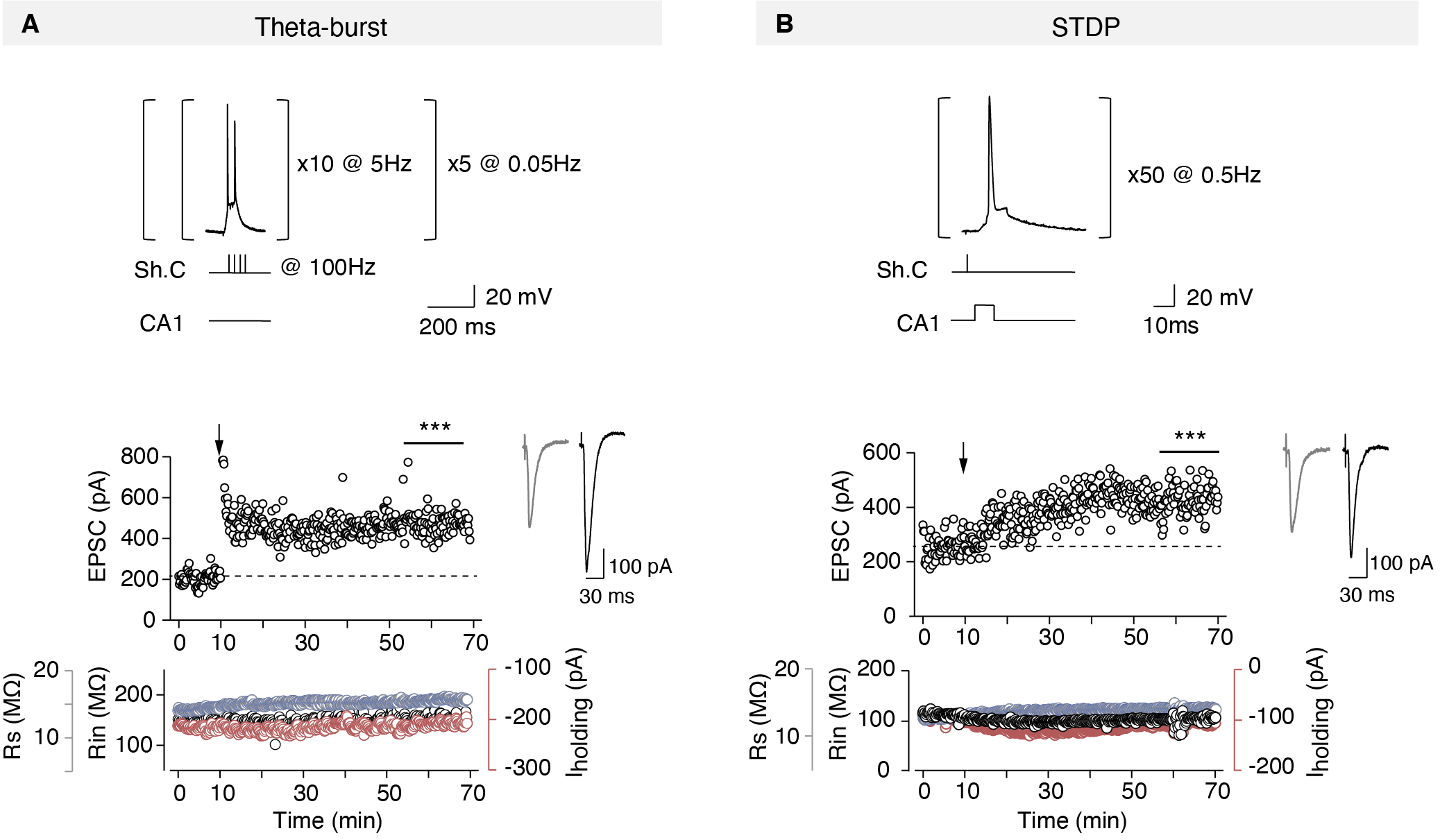

**Supplementary Figure 1.** **Representative 5-TBS-LTP and STDP-LTP (related to the main Figure 1).**

(**A**) Example of LTP induced by 5-TBS (baseline: 205±4pA, increased by 127%, to 446±6pA, one hour after pairings.). Bottom, time course of Ri (baseline: 145±1MΩ and 50-60 min after pairings: 154±1MΩ; change of 6%), Rs (baseline: 14.14±0.03MΩ and 50-60 min after pairings: 16.31±0.02MΩ; change of 15%) and holding current I_holding_ (baseline: -217±1pA and 50-60 min after pairings: -208±4pA; change of -4%). (**B**) Example of STDP-LTP induced by 50 pre-post pairings (Δt_STDP_=-8.7±0.4ms) (the mean baseline EPSC amplitude was 257±5pA before pairings and was increased by ~67% to 430±6pA one hour after pairings). Bottom, time course of Ri (baseline: 110±1MΩ and 50-60 min after pairings: 101±1MΩ; change of -8%), Rs (baseline: 12.72±0.02MΩ and 50-60 min after pairings: 14.05±0.03MΩ; change of 10%) and holding current I_holding_ (baseline: -93±1pA and 50-60 min after pairings: -109±1pA; change of 18%).

Insets correspond to the average EPSC amplitude during baseline (grey traces) and the last 10 min of recording after STDP pairings (red traces). Statistics (student *t*-test, first *vs* last 10 min of recording): *** *p*<0.001.

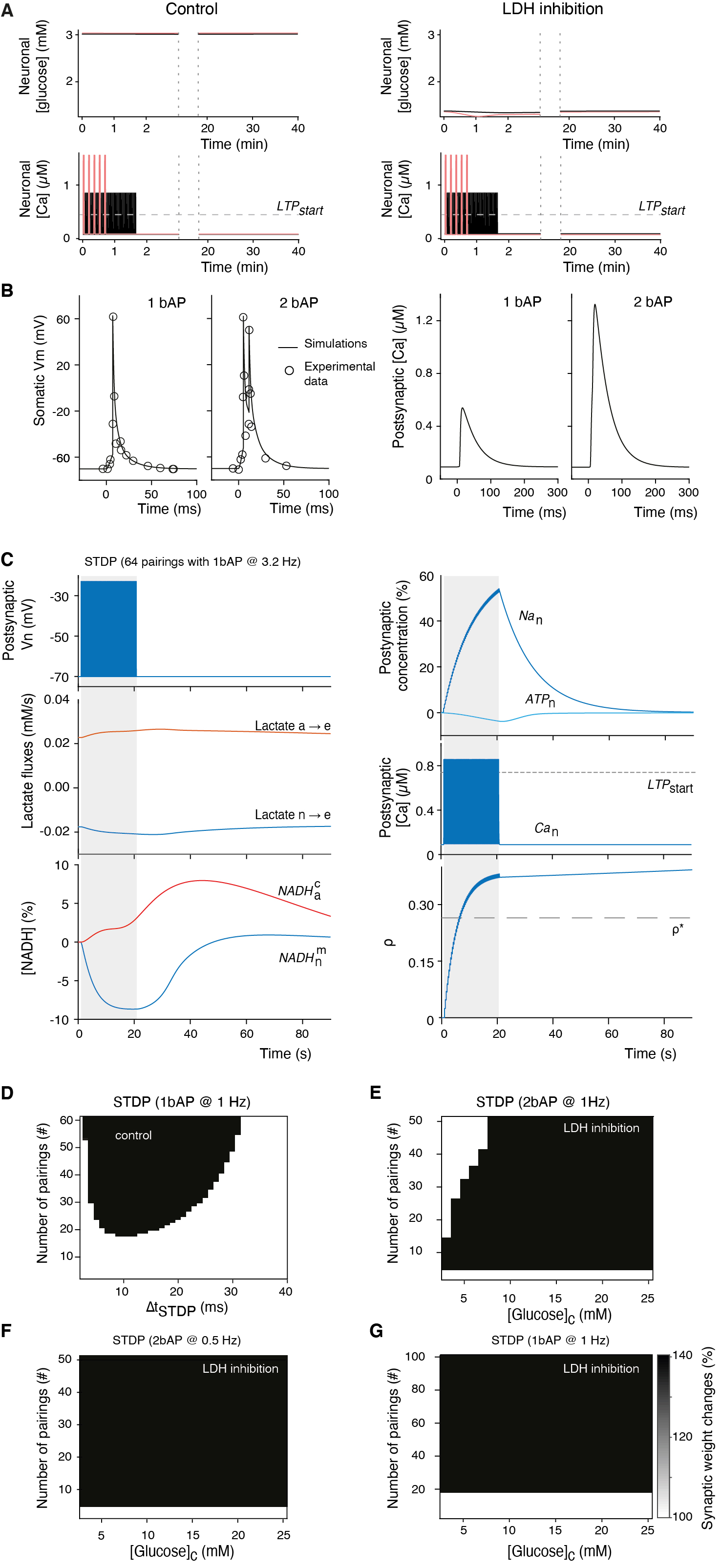

**Supplementary Figure 2.** **Model calibration (related to the main Figure 2).**

(**A**) Kinetics of neuronal glucose and calcium with 5-TBS (pink) or light STDP (50 pairings at 0.5Hz, black). (**B**) Experimental measurements (circles) and model prediction (line) of the somatic potential with 1 bAP or 2 bAP-stimulations and spike timing 10ms (*left*), together with the resulting calcium transients (*right*). (**C**) After calibration, the glia-neuron lactate shuttle part of the model reproduces experimental data in (*23*) where a 20 sec electrical stimulation of the neurons and astrocytes triggered a decay of neuronal mitochondrial NADH (${NADH}_{n}^{m}$) by 10% peaking around the end of the stimulation and converging back to baseline 15-20 min afterwards and a delayed overshoot of astrocytic cytosolic NADH (${NADH}_{a}^{c}$). (**D**) Prediction of synaptic plasticity by the model when the number of pairings and the spike timing vary in STDP (1 bAP at 1Hz) in control conditions. With the color code used, white means no plasticity, while the black zones correspond to LTP (**E-H**) explore the output of the model with LDH inhibited (oxamate), for STDP with 2 bAPs at 1Hz, 2 bAPs at 0.5Hz, and 1 bAP at 1 Hz, respectively, as a function of the number of pairings and the concentration of glucose in the bath solution.

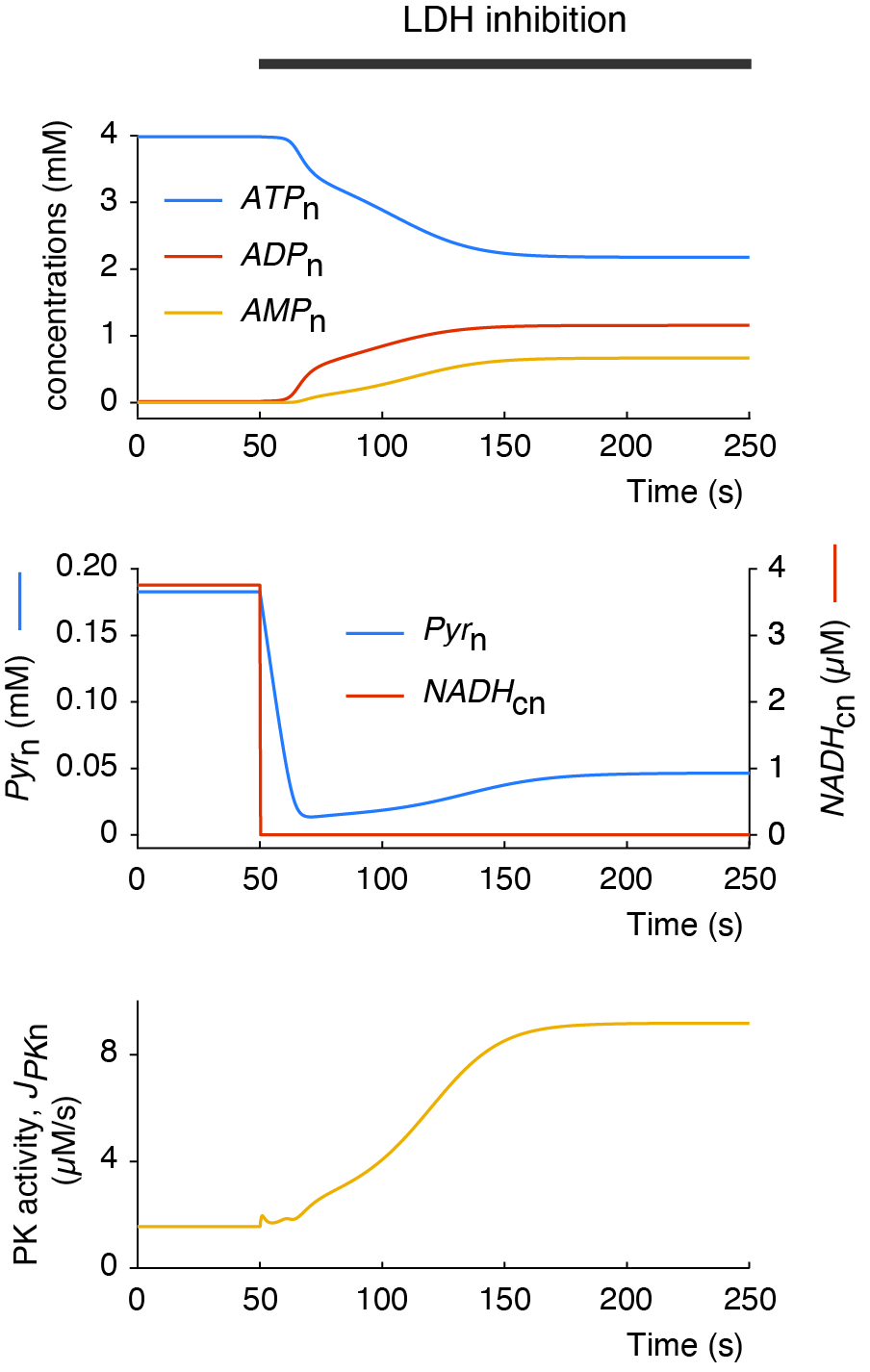

**Supplementary Figure 3. Model prediction of the effects of LDH inhibition on neuronal ATP levels (related to the main Figure 2).**

The model is initiated at time *t*=0 in control conditions. In control (*t*< 50s), ATP makes up roughly the totality of the 4 mM total adenosine phosphate concentration (*top*), the stationary level of pyruvate (*Pyr*_n_) and cytosolic NADH (*NADH*_cn_) in the neuron is large (*middle*), and neuronal glycolysis is low, as witnessed by the low activity of pyruvate kinase, PK (*bottom*). Oxamate addition in the neuron cytosol at *t*=50 sec rapidly switches the neuron to an oxidized redox state with a close to total depletion of cytosolic NADH. As a result, the ATP level drops well below 4 mM. Oxamate addition also results in neuronal glycolysis, as illustrated by the increase of pyruvate kinase activity. This restores significant levels of neuronal pyruvate and stabilizes ATP to roughly 50% of the total adenosine phosphate, *i.e.* slightly above 2 mM.

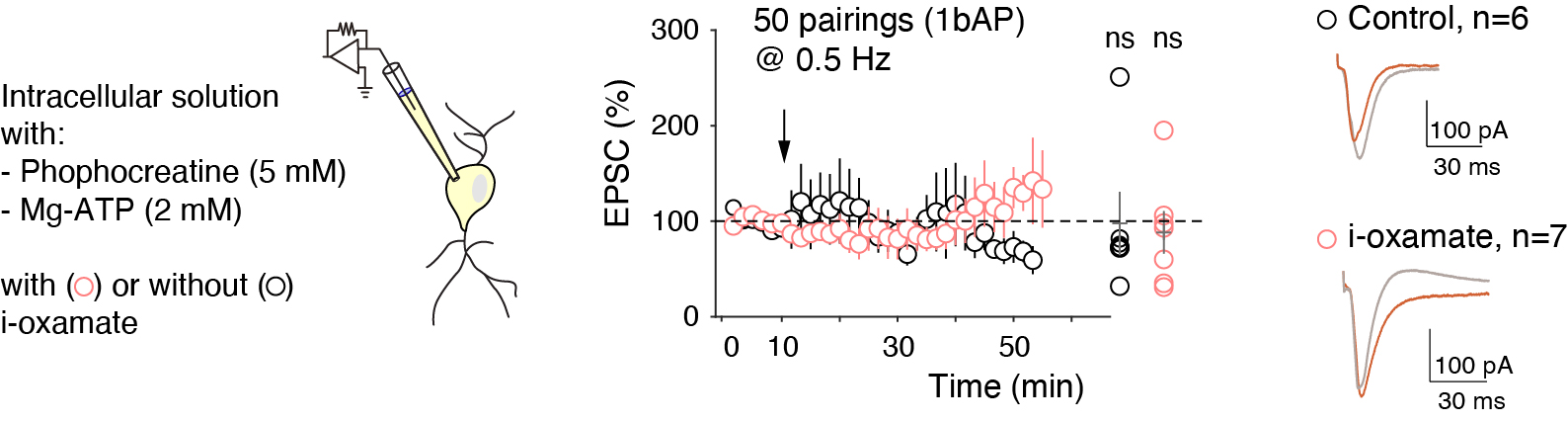

**Supplementary Figure 4. STDP is prevented with low intracellular ATP (2 mM) and phosphocreatine (5 mM) (related to the main Figure 2).**

Averaged time-course of the synaptic weight with EPSC amplitude 45-55 min after STDP. When intracellular solution contained 2 mM of ATP and 5 mM phosphocreatine, LTP was not observed after STDP protocol in control or i-oxamate conditions. Only one significant LTP could be induced in control and i-oxamate conditions out of 6 and 7 cells, respectively.

All data: mean±SEM. ns: not significant by two tailed *t*-test.

See Table S3 for detailed data and statistics.

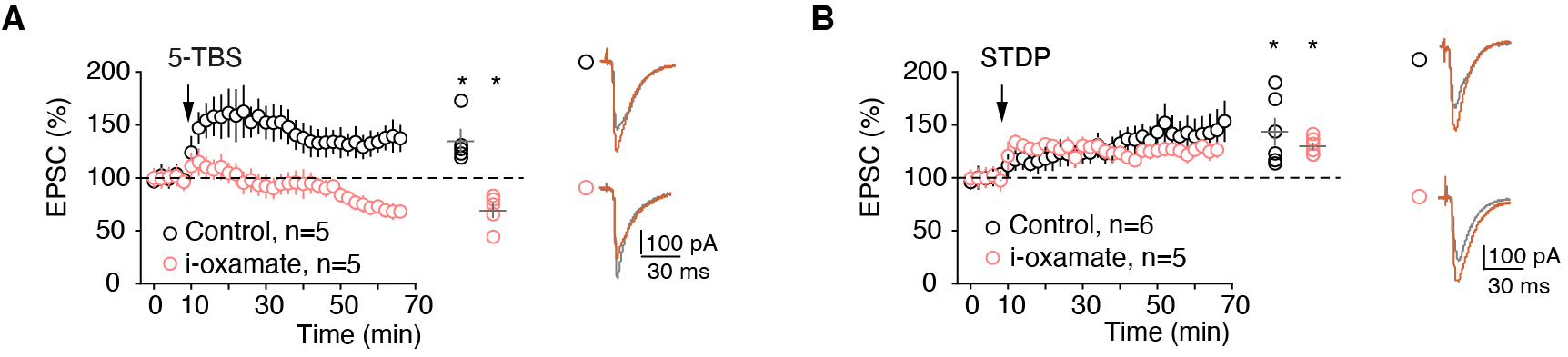

**Supplementary Figure 5. LDH inhibition prevents 5-TBS-LTP and left unaffected STDP-LTP in adult mice.**

Averaged time-course of the synaptic weight with EPSC amplitude 50-60 min after 5-TBS or STDP in adult mice. Intracellular inhibition of LDH with i-oxamate show distinct effects on 5-TBS and STDP expression since 5-TBS did not induce plasticity whereas STDP triggered a potent LTP.

Representative traces: 15 EPSCs averaged during baseline (grey) and 60 min (red) after protocols (arrows). All data: mean±SEM. *p<0.05; ns: not significant by two tailed *t*-test.

See Table S2 for detailed data and statistics.

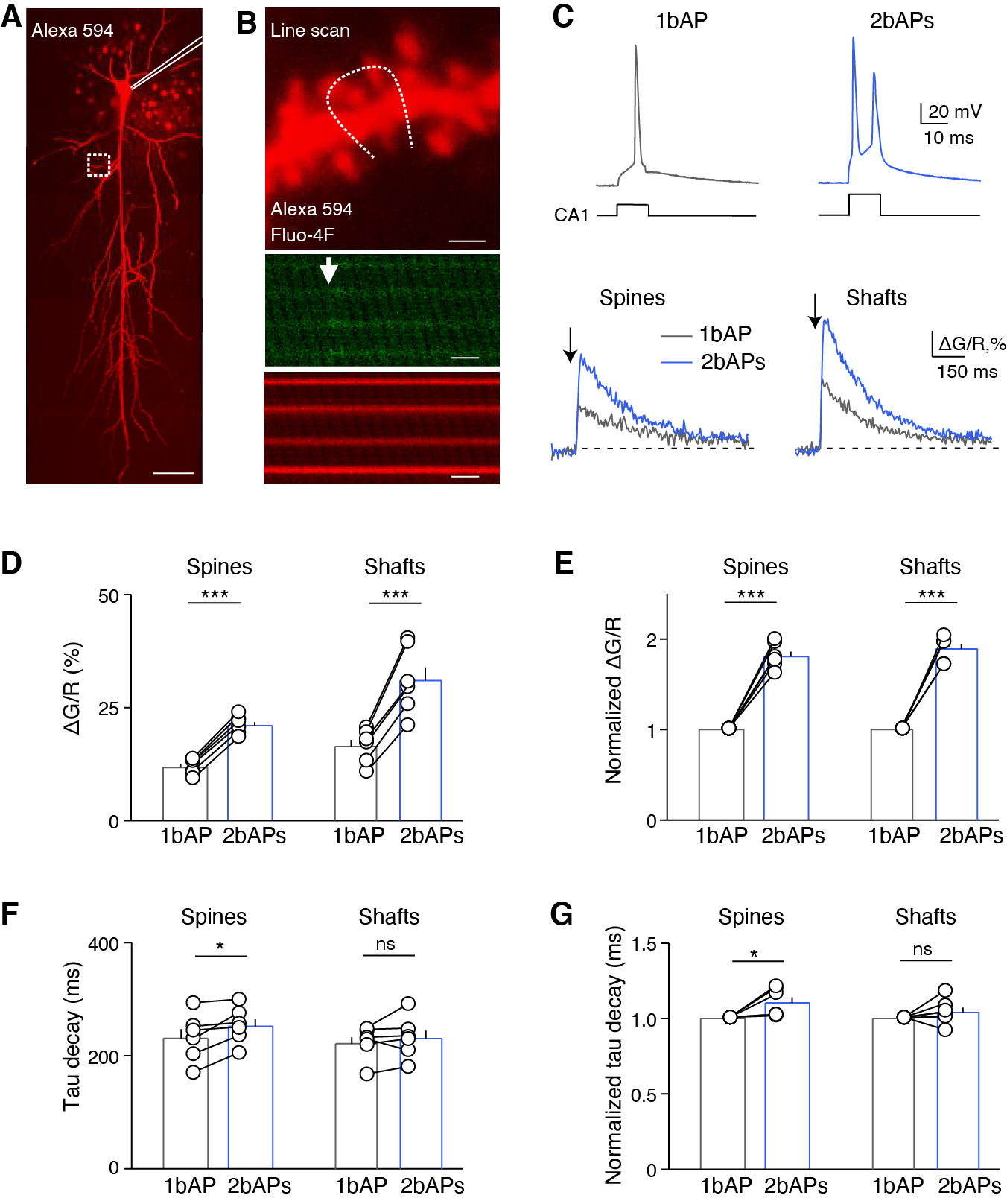

**Supplementary Figure 6.** **Calcium transients in dendritic spines and shafts upon single or two bAPs.**

(**A**) Combination of whole-cell recording of a CA1 pyramidal cell with two-photon imaging; the patch-clamp pipette is underlined in white and the dashed white square indicated the imaged dendritic area. The scanning areas had been selected on 50-150μm distance from soma. Scale bar: 30μm. (**B**) Line-scanning two-photon microscopy of Ca^2+^ transients in dendritic spines and adjacent shaft (top panel; scale bar: 2μm) filled with ratiometric indicators Fluo-4F (250μM; scale bar: 200ms) (middle panel) and Alexa Fluor 594 (50μM) (bottom panel; scale bar: 200ms). (**C**) A single or two bAPs were triggered in the recorded CA1 pyramidal cell by a postsynaptic current depolarization of 10 ms duration, and calcium transients were recorded in the dendritic spines and neighboring shafts. (**D** and **E**) Two bAPs induced larger increase of calcium in spines and shafts than a single bAP when evaluating ΔG/R (**D**) or the normalized ΔG/R (**E**). (**F** and **G**) Two bAPs induced a longer decay of the calcium transient in spines but not in shafts when compared to a single bAP, as estimated by the tau decay (**F**) and the normalized tau decay (**G**). Error bars represent the SEM. *: *p*<0.05; ***: *p*<0.001; ns: not significant.

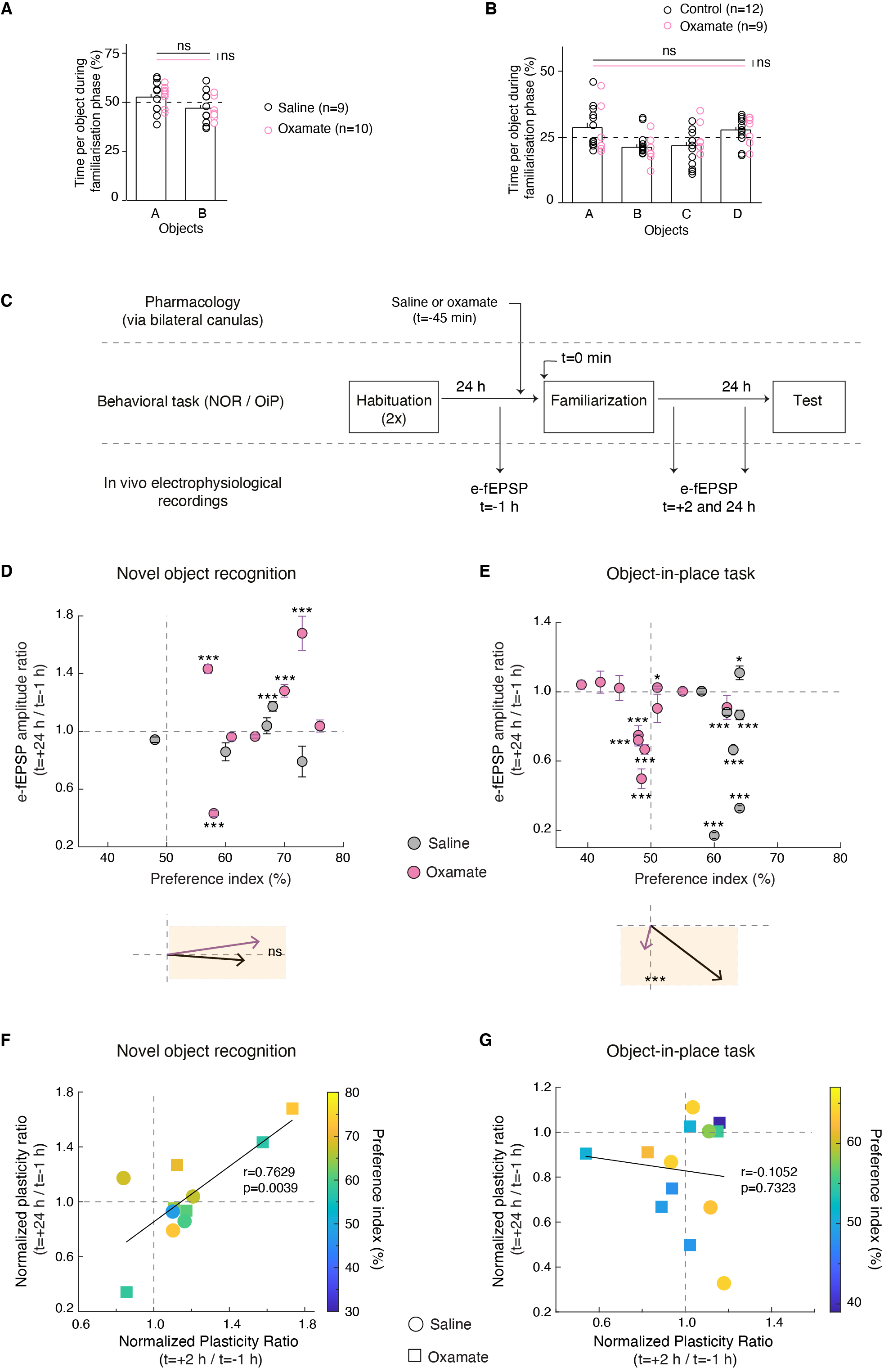

**Supplementary Figure 7.** ***In vivo*** **synaptic plasticity 24 h after test for NOR and OiP tasks (related to the main Figure 4).**

(**A**) Rats injected in CA1 with saline or oxamate solutions spent similar amount of time exploring A and B (saline: *p*=0.3750, oxamate: *p*=0.1661, saline *vs* oxamate: object A: *p*=0.5563, object B: *p*=0.5563, one-way ANOVA). (**B**) Saline- or oxamate-injected rats spent similar amount of time exploring A, B, C and D (saline: *p*=0.1342, oxamate: *p*=0.2178, saline vs oxamate: object A: *p*=0.8826, object B: *p*=0.1342, object C: *p*=0.2566, object D: *p*=0.6964, one-way ANOVA). A and B were common for both NOR and OiP tasks. (**C**) Behavioral experimental set-up for the NOR and OiP tasks and e-fEPSP recordings. (**D** and **E**) No difference for e-fEPSP amplitudes 24 hours after familiarization were observed between saline or oxamate-injected rats, with on average no plasticity for the NOR task (**D**) and a majority of LTD for the OiP task (**C**) (see average vectors in **D** and **E**). (**F** and **G**) Correlation between e-LFPs recorded 2 and 24 hours after familiarization in the NOR and OiP tasks, in relation with the preference index. NOR: significant positive correlation with the plasticity ratios decreasing from +2 to +24 hours, with larger LTPs remaining LTP at +24h, while smaller LTPs shifting into no plasticity or LTD (similar for both saline- and oxamate-injected rats). OiP: no correlation between plasticity at +2 and +24 hours with LTP at +2 hours turned either into no plasticity or LTD while the majority of LTD at +2 hours persisted in LTD at +24 hours.

All data are presented as mean±SD. *: p<0.05; **: p<0.01; ***: p<0.001; by two tailed *t*-test.

See Tables S4C and D for detailed data and statistics.

**Supplementary Tables**

**Supplementary Tables 1A and B (related to Figure 1):**

**A-**

|  | 5-TBS | | | STDP (50 pairings @ 0.5 Hz) | | |
| --- | --- | --- | --- | --- | --- | --- |
| Experimental conditions | EPSC amplitude % of baseline (n) | t-test,  *p* value | 2-way ANOVA, *p* value | EPSC amplitude % of baseline (n) | t-test,  p value | 2-way ANOVA, *p* value |
| Control | 152.4 ±13.8(14) | 0.0056 | <0.0001 | 138.3 ± 10.2 (8) | 0.0070 | <0.0001 |
| i-MK801 | 89.7 ± 6.4 (7) | 0.1562 | 0.0801 | 103.4 ± 7.0 (7) | 0.6484 | 0.4800 |
| DAB | 89.6 ± 10.5 (6) | 0.3691 | 0.0651 | 91.8 ± 9.5 (7) | 0.4213 | 0.0838 |
| DAB + e-lactate | 142.6 ± 17.0(6) | 0.0311 | 0.0032 | 159.3 ± 24.1(7) | 0.0383 | 0.0002 |
| i-oxamate | 91.7 ± 9.9 (9) | 0.4600 | 0.0643 | 134.8 ± 14.0 (7) | 0.0398 | 0.0008 |
| i-oxamate + i-pyruvate | 104.4 ± 13.3 (7) | 0.7548 | 0.5110 | - | - | - |
| i-oxamate + i-NADH | 158.5 ± 16.0 (6) | 0.0145 | <0.0001 | - | - | - |

**B-**

| Rescue experiments *vs.* corresponding control | T-test, *p* value |
| --- | --- |
| 5-TBS: control *vs.* e-DAB+ e-Lactate | 0.6735 |
| STDP (1 Hz, 50 pairings): control *vs.* e-DAB+ e-Lactate  5-TBS: i-oxamate + i-NADH *vs.* control | 0.4155  0.7995 |

**Supplementary Table 2. Parameter values of the mathematical model**

| **Parameters** | **Description** | **Value** | **Unit** |
| --- | --- | --- | --- |
| **Parameters estimated from our experimental data** | | | |
| $g_{L}$ | Postsynaptic leak conductance | 0.50 | mS/cm^2^ |
| $\tau_{\text{step}}$ | Postsynaptic depolarization current time constant | 15 | ms |
| $AT$ | Attenuation factor at the synapse | 0.34 | - |
| ${DP}_{\text{max}}$ | Amplitude of the postsynaptic depolarization | 1.50 (STDP 1bAP)  2.85 (STDP 2bAP)  0 (TBS) | mA/cm^2^ |
| ${DP}_{\text{dur}}$ | Duration of the postsynaptic depolarization | 10 (STDP)  0 (TBS) | ms |
| ${AP}_{\text{amp}}$ | Amplitude of the bAP at the soma | 120 (STDP 1bAP)  114 (STDP 2bAP)  130 (TBS) | mV |
| $\delta_{1}$ | Delay between depolarization onset and first bAP | 7 (STDP 1bAP)  5.5 (STDP 2bAP) | ms |
| $\delta_{2}$ | Delay between first and second bAP | 6.3 (STDP 2 bAP) | ms |
| $\alpha$ | Additional attenuation of the second bAP | 0.65 (STDP 2 bAP) | - |
| $\tau_{Ca}$ | Time scale of cytosolic Ca dynamics | 40 | ms |
| $\tau_{\rho}$ | Time scale of the internal state variable | 103.15 | s |
| $\rho^{*}$ | Unstable middle point of the bistable | 0.257 | - |
| ${LTD}_{\text{max}}$ | Maximal amplitude of the LTD | 70 | - |
| ${LTD}_{\text{start}}$ | Ca-threshold for the LTD | 0.45 | μM |
| ${LTP}_{\text{max}}$ | Maximal amplitude of the LTP | 120 | - |
| ${LTP}_{\text{start}}$ | Ca-threshold for the LTP | 0.74 | μM |
| $M_{\text{max}}$ | Max. amplitude of the ATP-gated depotentiation | 30 | - |
| ${ATP}_{\text{Thr}}$ | ATP-threshold of the ATP-gated depotentiation | 1.87 | mM |
| $\beta_{\rho}$ | Final synaptic weight increase in LTP | 0.405 (STDP 1baP)  0.405 (STDP 2baP)  0.550 (TBS) | - |
| $\tau_{AMPA1}$ | AMPA conductance time constant 1 | 9.6 | ms |
| $\tau_{AMPA2}$ | AMPA conductance time constant 2 | 7.0 | ms |
| $g_{AMPA\text{max}}$ | Max. AMPA conductance density | 0.13 | mS/cm^2^ |
| $\tau_{NMDA1}$ | NMDA conductance time constant 1 | 60.0 | ms |
| $\tau_{NMDA2}$ | NMDA conductance time constant 2 | 1.0 | ms |
| $g_{NMDA\text{max}}$ | Maximal NMDA conductance density | 4.64×10^-4^ | mS/cm^2^ |
| $g_{CaL\text{max}}$ | Maximal CaL conductance density | 0.0849 | mS/cm^2^ |
| $g_{Na\text{max}}$ | Maximal conductance density of VGSCs | 11.5 | mS/cm^2^ |
| **Fixed parameters and constants** | | | |
| $\left[ \text{Mg}^{2+} \right]$ | Magnesium concentration | 1.0 | mM |
| ${GLC}_{c}$ | Glucose concentration in the bath solution in control | 5.0 | mM |
| $E_{Ca}$ | Nernst Potential for Ca^2+^ | 54^1^ | mV |
| $E_{NMDA}$ | Reversal potential NMDA receptors | 0 | mV |
| $E_{L}$ | Reversal potential leak current | -70 | mV |
| $R$ | Gas constant | 8.3145 | J/mol/K |
| $T$ | Temperature | 310 | K |
| $F$ | Faraday Constant | 96 485.3 | C/mol |
| **Fixed parameters** taken from (*19*) with no change^2^ | | | |
| $S_{m}v_{x}$ | Surface/vol. ratio of compartment $x=\left\{ a,n \right\}$ | 2.5×10^4^ | cm^-1^ |
| $E_{AMPA}$ | Reversal potential AMPA receptors | 0 | mV |
| $C_{m}$ | Membrane Capacitance postsynaptic compartment | 10^-3^ | mF/cm^2^ |
| ${Na}_{e}$ | Sodium extracellular concentration | 150 | mM |
| $g_{Nax}$ | Na leak conductance in the postsynaptic compartment | 0.0136 (*n*)  0.0061 (*a*) | mS/cm^2^ |
| $V_{a}$ | Membrane voltage of the astrocyte | -70 | mV |
| $k_{\text{pump}x}$ | Max. rate Na-K-ATPases | 2.2×10^-6^ (*n*)  4.5×10^-7^ (*a*) | cm/mM/s |
| $K_{\text{Mpump}}$ | Affinity constant for ATP of Na-K-ATPases | 0.5 | mM |
| $K_{tg}$ | Affinity constant of glucose transporters | 8 | mM |
| ${Tg}_{xy}$ | Constant of glucose transport between compartments *x* and *y* | 0.0016 (*ca*)  0.0410 (*en*)  0.1470 (*ea*)  0.2390 (*ce*) | mM/s |
| $k_{HKPFKx}$ | Maximal rate combined HK-PFK | 0.0504 (*n*)  0.185 (*a*) | s^-1^ |
| $K_{g}$ | Affinity for glucose HKPFK | 0.05 | mM |
| $k_{PGKx}$ | Maximal rate PGK | 3.97 (*n*)  135.2 (*a*) | mM^-1^s^-1^ |
| *N* | Sum of NADH plus NAD^+^ concentrations in control | 0.212 | mM |
| *A* | Total adenine nucleotide concentration | 4.0 (*n*)  2.212 (*a*) | mM |
| *C* | Creatine plus phosphocreatine concentration | 10 | mM |
| $q_{AK}$ | Adenylate kinase equilibrium constant | 0.92 | mM |
| $k_{PKx}$ | Max. rate pyruvate kinase | 36.7 (*n*)  401.7 (*a*) | mM^-1^s^-1^ |
| $k_{LDH\text{on}x}$ | Forward rate constant of LDH | 72.3 (*n*)  1.59 (*a*) | mM^-1^s^-1^ |
| $k_{LDH\text{off}x}$ | Backward rate constant of LDH | 0.720 (*n*)  0.071 (*a*) | mM^-1^s^-1^ |
| ${LAC}_{c}$ | Lactate concentration in the reservoir | 0.55 | mM |
| ${Tl}_{xy}$ | Constant of lactate transport between compartments *x* and *y* | 24.3 (*ne*)  106.1 (*ae*)  0.00243 (*ac*)  0.25 (*ec*) | mM/s |
| $K_{tlxy}$ | Affinity constant of lactate transporters between *x* and *y* | 0.74 (*ne*)  3.50 (*ae*)  1.00 (*ac*)  1.00 (*ec*) | mM |
| $v_{\text{mitoin}x}$ | Maximal rate of the TCA cycle | 0.1303 (*n*)  5.7 (*a*) | mM/s |
| $K_{Mmito}$ | Affinity constant TCA cycle for pyruvate | 0.04 | mM |
| $K_{MNADx}$ | Affinity constant TCA cycle for NAD | 0.409 (*n*)  40.3 (*a*) | mM |
| $v_{\text{mitoout}x}$ | Maximal rate of the electron transport chain (ETC) | 0.164 (*n*)  0.064 (*a*) | mM/s |
| $K_{O2mito}$ | Affinity constant of ETC for O_2_ | 0.001 | mM |
| $K_{MADPx}$ | Affinity constant of ETC for ADP | 3.410 (*n*)  0.483 (*a*) | μM |
| $K_{MNADHx}$ | Affinity constant of ETC for NADH | 44.4(*n*)  26.9 (*a*) | μM |
| $T_{NADHx}$ | Maximal rate for NADH shuttling | 10330 (*n*)  150 (*a*) | mM/s |
| $M_{cytox}$ | Constant for shuttling from cytosol | 4.9×10^-8^ (*n*)  2.5×10^-4^ (*a*) | - |
| $M_{mitox}$ | Constant for shuttling from mitochondria | 3.93×10^5^ (*n*) 1.06×10^4^ (*a*) | - |
| $k_{\text{CKon}x}$ | Forward rate of creatine kinase | 0.0433 (*n*)  0.00135 (*a*) | mM^-1^s^-1^ |
| $k_{\text{CKoff}x}$ | Backward rate of creatine kinase | 2.8×10^-4^ (*n*)  10^-5^ (*a*) | mM^-1^s^-1^ |
| ${0_{2}}_{c}$ | O_2_ concentration in the bath solution | 7 | mM |
| ${{PS}_{cap}}/{V_{x}}$ | O_2_ transport constant from the reservoir | 1.66 (*n*)  0.87 (*a*) | s^-1^ |
| $K_{O2}$ | O_2_ exchange constant | 0.0361 | mM |
| $Hb.OP$ | O_2_ exchange constant | 8.6 | mM |
| $n_{h}$ | O_2_ exchange Hill number | 2.73 | - |
| $v_{x}$ | Relative volumes | 0.45 (*n*)  0.25 (*a*)  0.20 (*e*) | - |
| $\xi$ | Relative mitochondria volume | 0.07 | - |
| $J_{ATPasesx}$ | ATPase activity of non-Na-K-ATPases | 0.1695 (*n*)  0.1404 (*a*) | mM/s |
| $J_{pumpa0}$ | Basal Na-K-ATPase activity | 0 (*n*)  0. 0687 (*a*) | mM/s |

^1^: after compensation for the slight K^+^ permeability of CaL channels, see (*41*).

^2^: *n* is for the postsynaptic neuronal compartment, *a* for the astrocytic one, *e* for the extracellular pericellular volume and *c* for constant reservoir concentrations.

**Supplementary Tables 3A and B (related to Figures 2 & 3, Figures S4 & S5):**

**A-**

| Experimental conditions | 5-TBS  except for ^(2)^ | | | STDP (50 pairings @ 0.5 Hz)  except for ^(1, 4 and 5)^ | | |
| --- | --- | --- | --- | --- | --- | --- |
|  | EPSC amplitude % of baseline (n) | t-test,  *p* value | 2-way ANOVA, *p* value | EPSC amplitude % of baseline (n) | t-test,  p value | 2-way ANOVA, *p* value |
| Control, 25mM glucose | 169.7 ± 25.7 (8) | 0.0299 | <0.0001 | - | - | - |
| i-oxamate, 25mM glucose | 202.8 ± 20.9 (6) | 0.0044 | <0.0001 | - | - | - |
| i-Mannoheptulose | 134.6 ± 16.8 (7) | 0.0230 | 0.0013 | 176.7 ± 24.7 (7) | 0.0210 | <0.0001 |
| Control^(1)^ | - | - | - | 151.1±16.8 (8) | 0.0191 | <0.0001 |
| i-mannoheptulose^(1)^ | - | - | - | 176.3±29.9 (6) | 0.0436 | 0.0003 |
| i-Mannoheptulose  + i-oxamate^(1)^ | - | - | - | 86.6±19.9 (6) | 0.5297 | 0.2856 |
| i-oxamate^(1)^ | - | - | - | 151.1±16.8 (8) | 0.0191 | <0.0001 |
| Control^(2)^ | 209.9±33.9 (6) | 0.0229 | <0.0001 | - | - | - |
| i-oxamate^(2)^ | 162.4±15.8 (6) | 0.0109 | <0.0001 | - | - | - |
| i-oxamate^(3)^ | - | - | - | 80.8±9.4 (6) | 0.0956 | 0.0005 |
| i-oxamate+i-NADH^(3)^ | - | - | - | 119.4±4.7 (7) | 0.0061 | 0.0004 |
| i-oxamate^(4)^ | - | - | - | 228.0±38.8 (6) | 0.0216 | <0.0001 |
| Control^(5)^ | - | - | - | 123.1±8.6 (7) | 0.0369 | 0.0117 |
| i-oxamate^(5)^ | - | - | - | 129.0±9.8 (6) | 0.0309 | <0.0001 |
| Control ^(6)^ | - | - | - | 97.7±31.5 (6) | 0.9439 | 0.4643 |
| i-oxamate ^(6)^  Control (mice) ^(1)^  i-oxamate (mice) ^(1)^ | -  131.9±11.0 (5)  70.1±6.8 (5) | -  0.0441  0.0121 | -  0.0002  <0.0001 | 88.3±21.1 (7)  141.1±14.7 (6)  125.8±3.4 (5) | 0.5992  0.0454  0.0024 | 0.0001  <0.0001  <0.0001 |

^(1)^ STDP with 100 pairings at 1 Hz

^(2)^ 1-TBS

^(3)^ STDP with 50 pairings (2bAPs) at 1 Hz

^(4)^ STDP with 25 pairings (2Aps) at 1 Hz

^(5)^ STDP with 25 pairings (2bAPs) at 0.5 Hz

^(6)^ intracellular solution with 2 mM ATP and 5 mM phosphocreatine

**B-**

| Experimental conditions | T-test, *p* value |
| --- | --- |
| STDP (1 Hz, 100 pairings, 1 bAP): control vs. i-oxamate | 0.9836 |
| STDP (1 Hz, 100 pairings): control vs. i-oxamate+i-lactate | 0.7095 |
| STDP (1 Hz, 100 pairings): control vs. i-oxamate | 0.9837 |
| STDP (1 Hz, 100 pairings): i-oxamate vs. i-oxamate+i-lactate | 0.5901 |

**Supplementary Tables 4A-D (related to Figure 4 and Figure S7):**

**A-**

| NOR  task | Saline-injected rats | | Oxamate-injected rats | | Saline vs.  oxamate |
| --- | --- | --- | --- | --- | --- |
|  | Time per object  (N) vs (F) /  Preference index (n) | t-test,  *p* value | Time per object / Preference index (n) | t-test,  *p* value | t-test,  *p* value |
| All | (N) 66.6±3.1%  (F) 33.4±3.1% /  66.6±3.1 (9) | 0.0007 | (N) 67.6±3.2%  (F) 32.4±3.2% /  67.6±3.2 (10) | 0.0003 | 0.8126 |
| AA-AB | (N) 71.8±2.9%  (F) 28.2±2.9% /  71.8±2.9 (4) | 0.0051 | (N) 73.0±4.4%  (F) 27.0±4.4% /  73.0±4.5 (5) | 0.0063 | 0.8309 |
| BB-BA | (N) 62.4±4.4%  (F) 47.6±4.4% /  62.4±4.4 (5) | 0.0475 | (N) 62.3±3.3%  (F) 37.7±3.3% /  62.3±3.3 (5) | 0.0213 | 0.9792 |
| AA-AB ^(1)^ | (N) 63.4±4.3%  (F) 36.6±4.3% /  63.4±4.3 (5) | 0.0370 | (N) 65.9±2.8%  (F) 47.6±2.8% /  65.9±2.8 (7) | 0.0013 | 0.7617 |

^(1)^: rats with chronic recording and stimulation electrodes (e-fEPSPs), with bilateral cannulas.

(N) and (F): novel and familiar, respectively, objects.

**B-**

| OiP  task | Saline-injected rats | | Oxamate-injected rats | | Saline vs.  oxamate |
| --- | --- | --- | --- | --- | --- |
|  | Time per object  (N) vs (F) /  Preference index (n) | t-test,  *p* value | Time per object / Preference index (n) | t-test,  *p* value | t-test,  *p* value |
| All | (N) 63.7±2.3%  (F) 36.3±2.3% /  63.7±2.3 (12) | <0.0001 | (N) 46.4±3.0%  (F) 53.7±3.0% /  46.4±3.0 (9) | 0.2500 | 0.0002 |
| AC | (N) 68.6±3.2%  (F) 31.4±3.2% /  68.6±3.2 (6) | 0.0020 | (N) 53.0±4.7%  (F) 46.9±4.7% /  53.0±4.7 (4) | 0.5603 | 0.0209 |
| BD | (N) 58.9±2.0%  (F) 41.1±2.0% /  58.9±2.0 (6) | 0.0068 | (N) 41.0±1.3%  (F) 59.0±1.3% /  41.0±1.3 (5) | 0.0026 | <0.0001 |
| AC ^(1)^ | (N) 61.7±1.2%  (F) 38.3±1.2% /  61.7±1.2 (9) | <0.0001 | (N) 49.1±1.7%  (F) 50.9±1.7% /  49.1±1.7 (12) | 0.6239 | <0.0001 |

^(1)^: rats with chronic recording and stimulation electrodes (e-fEPSPs), with bilateral cannulas.

(N) and (F): novel and familiar, respectively, objects.

**C- e-fEPSP plasticity at 2 and 24 hours after familiarization phase in saline- and oxamate-injected rats in NOR and OiP tasks.**

| e-fEPSP  monitoring | Saline-injected rats | | Oxamate-injected rats | | Saline *vs.*  oxamate |
| --- | --- | --- | --- | --- | --- |
|  | *t* post-familiarization:  +2 h (n)  + 24 h (n) | t-test,  *p* value | *t* post-familiarization:  +2 h (n)  + 24 h (n) | t-test,  *p* value | t-test,  *p* value |
| NOR task | 108.2±6.4 (5)  96.1±6.7 (5) | 0.2686  0.5904 | 128.0±10.7 (7)  112.1±15.3 (7) | 0.0395  0.4570 | 0.1839  0.4210 |
| OiP task | 114.9±5.6 (7)  71.8±15.7 (7) | 0.0377  0.0778 | 93.7±6.3 (9)  87.3±6.6 (12) | 0.3447  0.0471 | 0.0287  0.2380 |

**D- Comparison of average vectors in saline- and oxamate-injected rats (MANCOVA)**

| Average vectors in saline- and oxamate-injected rats (MANCOVA) | Univariate test on the  e-fEPSP plasticity axis | | Univariate test on the  behavior axis | | Multivariate Test | |
| --- | --- | --- | --- | --- | --- | --- |
|  | F | *p* value | F | *p* value | F | *p* value |
| NOR task  +2h  +24h | 2.038  0.704 | 0.184  0.421 | 0.260  0.260 | 0.621  0.621 | 0.980  0.353 | 0.412  0.712 |
| OiP task  +2h  +24h | 5.94  1.50 | 0.029  0.238 | 16.43  28.82 | 0.01   < 0.001 | 14,7  13.9 | <0.001  <0.001 |
